# The innate immune DNA sensor cGAS is a negative regulator of DNA repair hence promotes genome instability and cell death

**DOI:** 10.1101/465401

**Authors:** Hui Jiang, Swarupa Panda, Xiaoyu Xue, Fengshan Liang, Patrick Sung, Nelson O. Gekara

## Abstract

Stringent regulation of DNA repair is essential for organismal integrity, but the mechanisms are not fully understood. Cyclic cGMP-AMP synthase (cGAS), the DNA sensor that alerts the innate immune system to the presence of foreign or damaged self-DNA in the cytoplasm is critical for the outcome of infections, inflammatory diseases and cancer. Besides this cytoplasmic function as an innate immune sensor, whether cGAS fulfills other biological roles remains unknown. Here we report that cGAS has a distinct role in the nucleus: it inhibits homologous recombination DNA repair (HR) thereby promoting genome instability and associated micronuclear generation and mitotic death. We show that cGAS-mediated inhibition of HR requires its DNA binding and oligomerization but not its catalytic activity or the downstream innate immune signaling events. Mechanistically, we show that cGAS impede RAD51-mediated DNA strand invasion, a key step in HR. These results uncover a new function of cGAS relevant for understanding its involvement in genome instability- associated disorders.

## INTRODUCTION

The DNA Damage Response (DDR) that senses threats to our genome, and the immune system - the inherent ability to sense and respond to infections, both function as surveillance systems essential for the preservation of organismal integrity. Emerging evidence indicates that these two systems are interdependent ^1^.

DDR constitutes a complex set of signalling pathways that mediate the elimination of DNA lesions, but could also trigger senescence or cell death to prevent the transmission of damaged genetic material ^2, 3^. Besides these classical outcomes of DDR, it is increasingly becoming apparent that DNA damage has a major impact on the immune system. DNA damage releases DNA fragments into the cytosol resulting in innate immune induction of cytokines such as type I interferons (IFN-Is) ^1^. These cytokines in turn play an important role in priming the immune system for enhanced antimicrobial ^1^ and antitumor immunity ^4-7^, but, if produced in excessive amounts, could lead to autoimmunity ^1, 8^ and tumorigenesis ^9-11^. Moreover, DNA damage-induced cytokines have also been found to be essential for classical DDR cellular outcomes including senescence ^12-15^.

Altogether, these recent findings highlighting the symbiotic partnership between DDR and the immune system have opened new avenues for understanding the biology of several disorders such as cancer and autoimmune diseases, caused by genome instability and immune defects ^10^. However, it remains unclear how these two biological systems are cross-regulated and what signalling molecules are involved.

From recent reports, there is a growing consensus that a key player linking DNA damage and immunity is the cyclic cGMP-AMP synthase (cGAS). cGAS surveys the cytoplasm for the presence of microbial DNA ^16, 17^, self-DNA from damaged chromatin ^1^, or DNA released from distressed mitochondria ^18^. Upon recognition of double stranded DNA (dsDNA), cGAS catalyzes the cyclization of ATP and GTP into the second messenger cyclic GMP–AMP (2′3′-cGAMP) ^16, 17, 19-24^. Subsequently, cGAMP binds to its adaptor Stimulator of Interferon Genes (STING) ^25^ leading to the activation of downstream transcription factors: Interferon regulatory factor 3 (IRF3) and nuclear factor kB (NF-kB) ^26, 27^, and these together, coordinate the induction IFN-Is and other inflammatory cytokines. Dysfunctions in the cGAS-STING pathway have been implicated in many disorders including infections, inflammatory diseases, neurodegeneration and cancer ^10, 28^.

Thus far, all the functions ascribed to cGAS are linked to its cytoplasmic role in innate immune system activation. Beyond this, whether cGAS also serves other biological functions is unknown.

DNA double-strand DNA breaks (DSB) are potentially highly deleterious lesions. If improperly repaired, DSB results in chromosomal deletions or translocations culminating in genome instability-associated disorders including tumorigenesis, accelerated aging and other diseases ^3^. For a healthy outcome, the DNA damage signaling is carefully calibrated to ensure timely removal of damaged DNA breaks, but if the DNA damage is excessive, to promote cell death to eradicate genetically defective cells ^29^. The regulatory molecules involved in this delicate balance are not fully known. In this study, we report our studies identifying a new regulator of DNA repair: cGAS. We demonstrate that aside from its function in the cytoplasm as an innate immune sensor, cGAS is also present in the nucleus where it acts as a negative regulator of homologous recombination-mediated DNA repair and thereby promotes micronuclear generation and mitotic catastrophic death of genetically stressed cells.

## RESULTS

### cGAS is presence in the nuclear and cytosolic compartments

cGAS is generally considered a cytosolic protein. However, while monitoring the subcellular localization of endogenous cGAS in bone morrow differentiating monocytes (BMDMos) or exogenously expressed GFP-tagged human cGAS (GFP-hcGAS) in HEK293 cells, we noted that cGAS is also abundant in the nucleus (**Figure 1 and Figure S1**). Curiously, and consistent with recent observations ^13^, when cells were enriched at G0/G1 cell cycle phase by serum starvation or by cell contact inhibition at high cell density, or at G1/early S phase by the DNA polymerase  inhibitor aphidicolin, cGAS showed an increase in the cytosol (**Figure 1 and Figure S1**). These data demonstrate that cGAS is constitutively present in the cytosol and nucleus and that the relative abundance of cGAS in these subcellular compartments can vary with the cell cycle. To better understand the cGAS features essential for its nuclear localization we analysed different cGAS mutants. The nuclear localization of the catalytic dead E225A/D227A mutant (GFP-hcGASΔcGAMP)^30^ and the oligomerization defective K394A mutant (GFP-hcGASΔOligo)^21^ were comparable to that of full length GFP-hcGAS. In contrast, DNA binding C396A/C397A mutant (GFP-hcGASΔDNA)^19, 31^ did not show a strong accumulation in the nucleus and was unaffected by changes in cycle phase (**Figure S2A**). On further analysis we found that nuclear cGAS was not within soluble nuclear fractions but mostly chromatin-bound (**Figure S2B**). Together with previous observations^13^, these findings indicated that the localization of cGAS in the nucleus is due to its binding to DNA following dissolution of the nuclear membrane during cell division. To interrogate this further, we asked how introduction a strong nuclear export or import signals (NES or NLS respectively) would impact cGAS localization. The NLS localized cGAS almost entirely into the nucleus. In contrast, the NES substantially increased hcGAS localization in the cytosol but did not eliminate its presence from the nucleus (**Figure S2C, D**). Taken together with the behavior of the cGASΔDNA mutant (**Figure S2A**), these results confirm that the localization and retention of cGAS in the nucleus is due to its avid binding to DNA and hence even a strong NES is not sufficient to not completely dislodge it from the nucleus.

**Figure 1.**
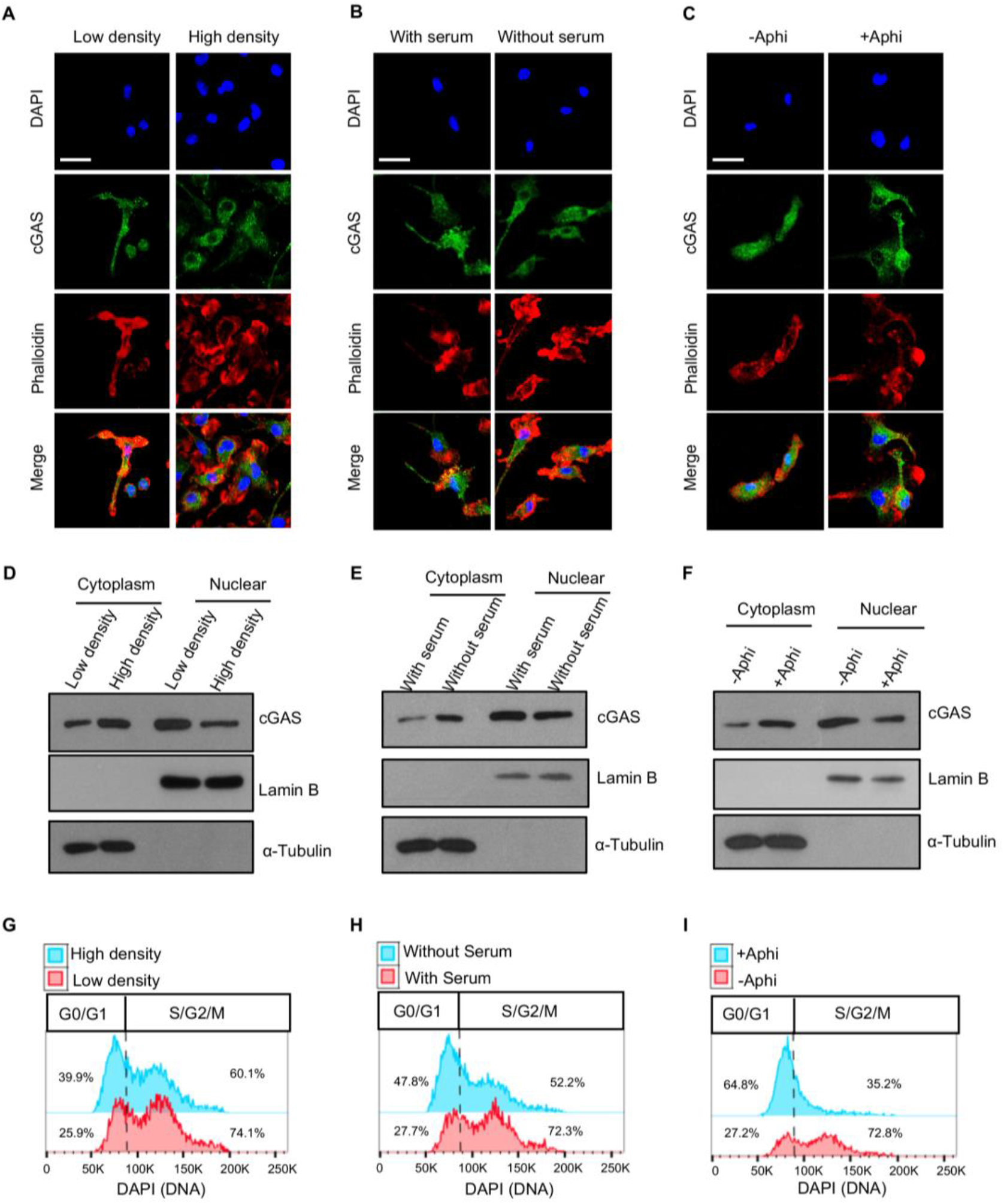
cGAS is constantly present in the cytosol and nucleus and is impacted by cell cycle. **(A-C)**, Immunofluorescence images of cGAS in the nucleus (DAPI) and cytosol (phalloidin) in BMDMos cultured at low/high density (**A**), with/without serum (**B**), or with/without Aphidicolin (Aphi) (**C**). Scale bar: 50 μm. (**D-F).** Immunoblot estimation of cGAS in nuclear/cytosolic subcellular fraction of BMDMos cultured under indicated conditions. Lamin B and α-Tubulin are nuclear and cytosolic markers respectively. (**G-I**) Flow cytometric analysis of cell cycle of BMDMos depicted in D-F. See also Figure S1.

### Nuclear cGAS promotes genome destabilization, micronuclei generation and mitotic death

We wished to determine the biological role of nuclear cGAS. Micronuclei are hallmarks of genome instability. Micronuclei arise following the mis-segregation of broken chromosomes during mitosis ^32, 33^ and have recently been proposed as platforms for cGAS-mediated innate immune activation following DNA damage ^6, 33, 34^. While studying the impact of cGAS in the cellular response to genotoxic stress, we noted that in response to γ-irradiation, HEK293 cells expressing GFP-hcGAS exhibit a higher incidence of micronuclei than cells expressing a GFP control containing a nuclear localization sequence (GFP-NLS) (**Figure. 2A, B**). As expected, GFP-hcGAS, but not GFP-NLS, restored *IFNB1* response to transfected DNA (**Figure. 2C**). This observation led us to hypothesize that the presence of cGAS in the nucleus and micronuclei generation were causally related. Hence we tested whether endogenous cGAS does promote micronuclei generation in BMDMos. To induce micronuclei generation, BMDMos were arrested at the G2 phase using the microtubule-depolymerizing agent nocodazole followed by γ-irradiation and release into mitosis (**Figure 2D**). BMDMos from WT mice exhibited more micronuclei compared to those from *cGAS^−^/-* mice (**Figure 2D-F**), demonstrating that in addition to innate immune activation, cGAS promotes genome destabilization.

**Figure 2.**
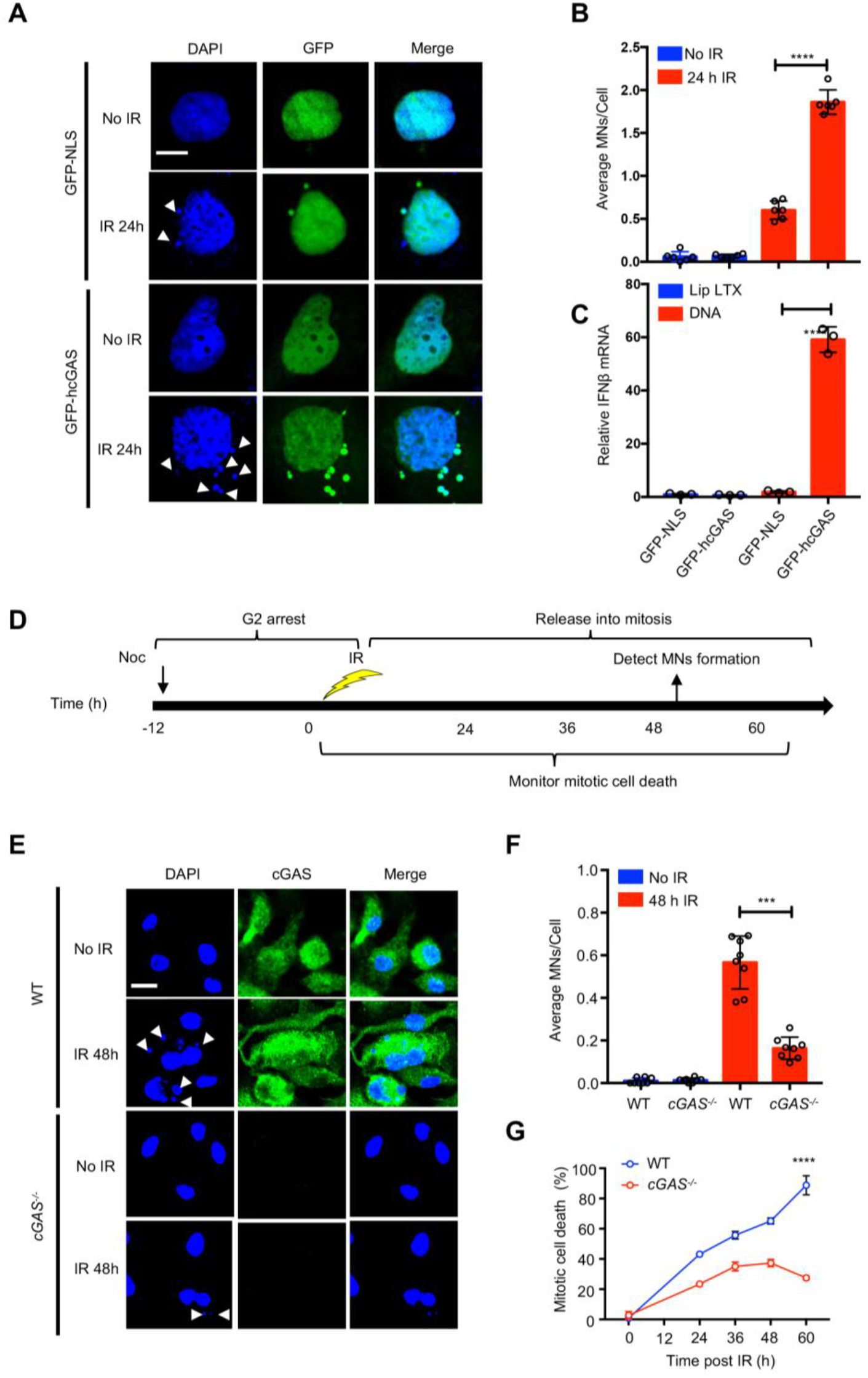
cGAS promotes micronuclei generation and mitotic cell death. **(A-B)**, Micronuclei (MN) form more frequently in HEK293 cells expressing GFP-hcGAS. (**A**) Micronuclei (indicated by arrow head) in GFPNLS- or GFP-hcGAS-expressing HEK293 cells before (0 h) or 24 h after γ-irradiation (IR; 10 Gy). Scale bar: 10 μm. (**B**) The average MNs/cell. (**C**), *IFNB1* response in HEK293 cells stimulated with transfected plasmid DNA. Mean ± SEM. of n=3 independent experiments; unpaired two-tailed Student’s t-test. **** P ≤ 0.0001. (**D**), Experimental outline for micronuclei generation and mitotic cell death in BMDMos. (**E**), Micronuclei (indicated by arrow head) and cGAS staining in WT and *cGAS^−^/-* BMDMos exposed to γ-irradiation (10 Gy). scale bar10 μm. (**F**), Average MNs/cell in corresponding representative images. MN graphs show Mean ± SEM independent experiments (n=3) representing eight different microscopic fields with over 200 cells; unpaired two-tailed Student’s t-test. *** P ≤ 0.001. (**G**), WT and *cGAS^−^/-* BMDMos arrested at G2, γ-irradiated (10 Gy) then evaluated for cell death at indicated time points after release into mitosis.

Mitotic catastrophe is a form of cell death, which, similar to micronuclei generation, occurs when damaged chromosomes mis-segregate during mitosis. Hence mitotic catastrophe is considered to be an important mechanism that prevents replication of genetically damaged cells ^29^. To further elucidate the biological relevance of nuclear cGAS, we tested the impact of cGAS on mitotic catastrophe. BMDMos from *cGAS^−^/-* mice were found to be less susceptible to mitotic catastrophic death compared to those from WT mice (**Figure 2G**). This demonstrates that by accelerating genomic destabilization, the concomitant chromosome mis-segregation and mitotic catastrophic death, cGAS contributes to the elimination of cells with severely damaged genomes.

### cGAS impedes DNA repair independently of STING signalling

Next we sought to determine whether cGAS contributes to genome instability by inhibiting DNA double strand break (DSB) repair. For that, we monitored DSB at different time points following γ-irradiation by comet and pulsed-field gel electrophoresis assays. We found that γ-irradiated BMDMos from *cGAS^−/−^* mice resolved DSB faster than those from WT mice (**Figure S3A, Figure 3A, B**), confirming that cGAS is a negative regulator of DSB repair. Curiously, while γ-irradiated BMDMos from *cGAS^−/−^* mice exhibited a faster resolution of DSB, those from *Sting^−/−^* mice were comparable to WT BMDMos in this regard (**Figure 3A, B**). Further, expression of GFP-hcGAS in the HEK293T cells that lack endogenous expression of both cGAS and STING ^17^ also impaired DSB repair in these cells (**Figure S3B**), but, as expected, failed to restore the *IFN-1* response (**Figure S3C**). Thus, in contrast to innate immune activation, cGAS-mediated inhibition of DSB repair is STING-independent.

**Figure 3.**
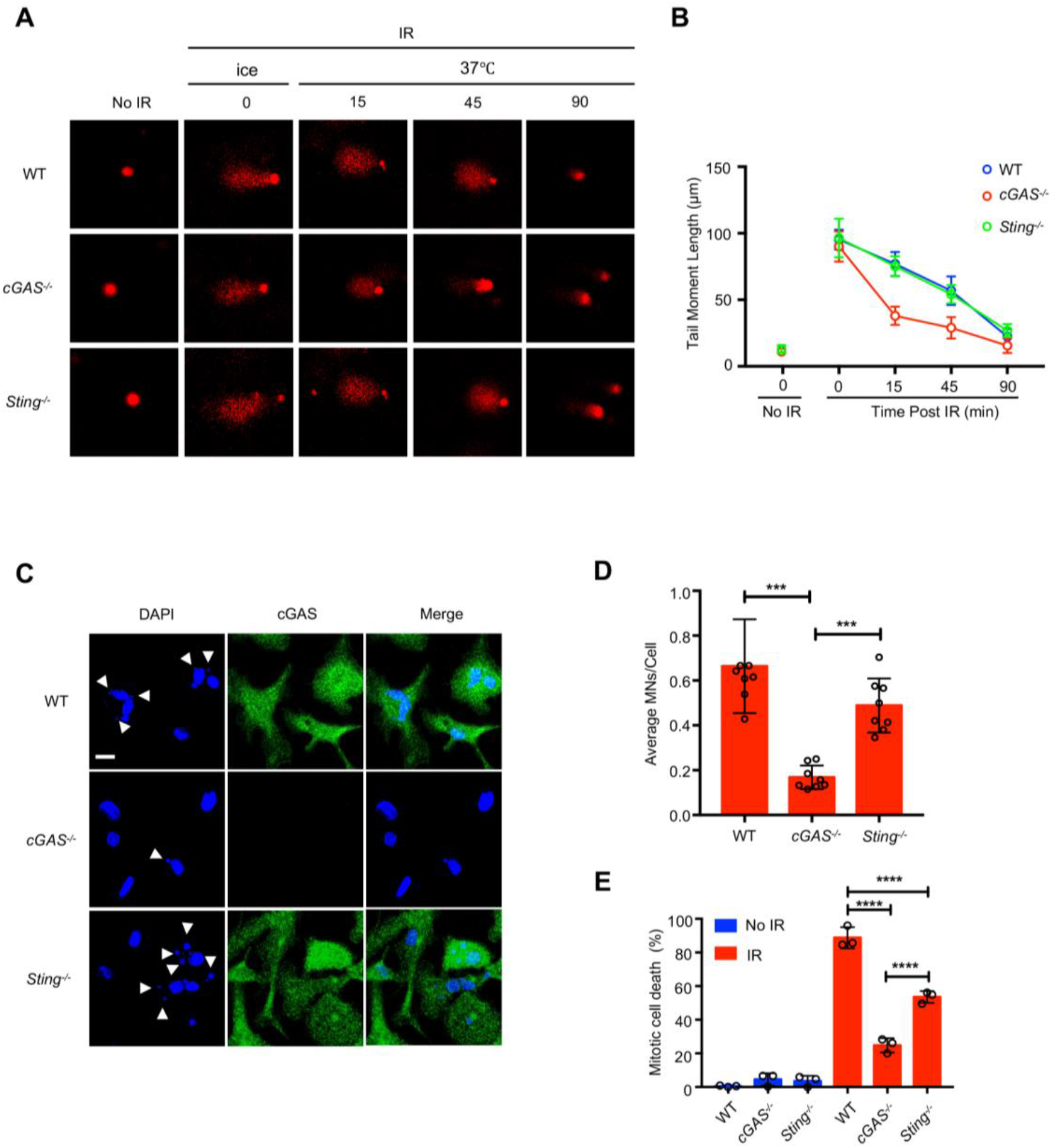
STING is dispensable for cGAS-mediated inhibition of DNA repair but boosts mitotic death. **(AB)**, BMDMos from WT and *Sting^−^/-* mice exhibit lower DNA repair efficiency than those from *cGAS^−^/-* mice. (**A**), Representative comet tails of WT, *cGAS^−^/-* and *Sting^−^/-* BMDMos exposed to γ-irradiation (IR: 10 Gy) on ice then incubated at 37°C for indicated duration. (**B**) Corresponding quantification of the comet tail moments from 20 different fields with *n* > 200 comets. Unpaired two-tailed Student’s t-test. *P ≤ 0.05, ***P ≤ 0.001. (**C-D**), cGAS promotes micronuclei generation in BMDMos. (D), Micronuclei (indicated by arrow head) and cGAS staining in WT, *cGAS^−^/-* and *Sting^−^/-* BMDMos exposed to γ-irradiation (10 Gy). Scale bar10 μm (**D**) Average MNs/cell in corresponding representative images. Bar graphs show mean values from eight different microscopic fields with over 200 cells. Mean ± SEM. of n=3 independent experiments; unpaired two-tailed Student’s t-test. *** P ≤ 0.001. (**E**), Mitotic cell death in WT, *cGAS^−^/-* and *Sting^−^/-* BMDMos.

To further interrogate how cGAS affects genome stability, we considered whether inhibition of DSB repair was mediated by cGAMP via a hitherto undefined STING-independent mechanism. However, treatment of HEK293 cells with cGAMP before or during γ-irradiation did not lead to increased fragmentation of genomic DNA (**Figure S3D**), but, as expected, activated STING-dependent Interferon Regulatory Factor (IRF3) (**Figure S3E**). Accordingly, the full length and catalytically dead GFP-hcGASΔcGAMP comparably inhibited DSB repair in HEK293 cells, as assessed by comet tail length (**Figure S3F, G**) and as expected, unlike GFP-hcGAS, GFP-hcGASΔcGAMP failed to restore the IFN-I response (**Figure S3H**).

Curiously, on further analysis we noted that although not essential for cGAS-mediated suppression of DNA repair, cGAMP-STING was partly required for optimal generation of micronuclei and induction mitotic cell death (**Figure 3C-D** and **Figure S3I, J**). This is consistent with previous findings indicating a role for interferons in mitotic stress^35, 36^ – an ingredient for mitotic death and micronuclei generation. Thus, although not primarily required for cGAS-mediated inhibition of DSB repair, STING-IFN-I signaling does contribute to secondary cellular outcomes of DNA damage such as micronuclei generation and mitotic death. Finally, when we compared hcGAS and mcGAS, we found them to inhibit DNA repair comparably (**Figure S3K, L**).

### cGAS attenuates DSB repair in a cell cycle and ATM-dependent manner

DSB repair occur via two major pathways: non-homologous end-joining (NHEJ) and homologous recombination (HR)^37, 38^. NHEJ is an error-prone repair pathway active throughout the cell cycle, and entails the ligation of DNA ends, often leading to deletion or insertional mutations ^37, 38^. HR is on the other hand an accurate repair process active in proliferating cells and occurs mainly during the S and G2 cell cycle phases wherein it engages the undamaged sister chromatid to template break repair to restore the original DNA sequence 37, 38.

To determine which of the two major DSB repair pathways is impeded by cGAS, we monitored γ-irradiation-induced DNA breaks under different conditions permissive or not permissive for either of these DNA repair pathways. We found that arresting cells at the G1/early S phase using Aphidicolin abolished the cGAS phenotype. In contrast, arresting cells at the G2 phase by nocodazole did not (**Figure 4A-D**). ATM and DNA-PKc are proximal kinases that play critical roles in HR and NHEJ, respectively. Inhibition of ATM abrogated the cGAS-associated increase in DNA breaks, while the DNA-PKc inhibitor had no effect (**Figure 4E-G, Figure S4A**), implying that cGAS mainly interferes with the ATM-dependent HR pathway.

**Fig. 4.**
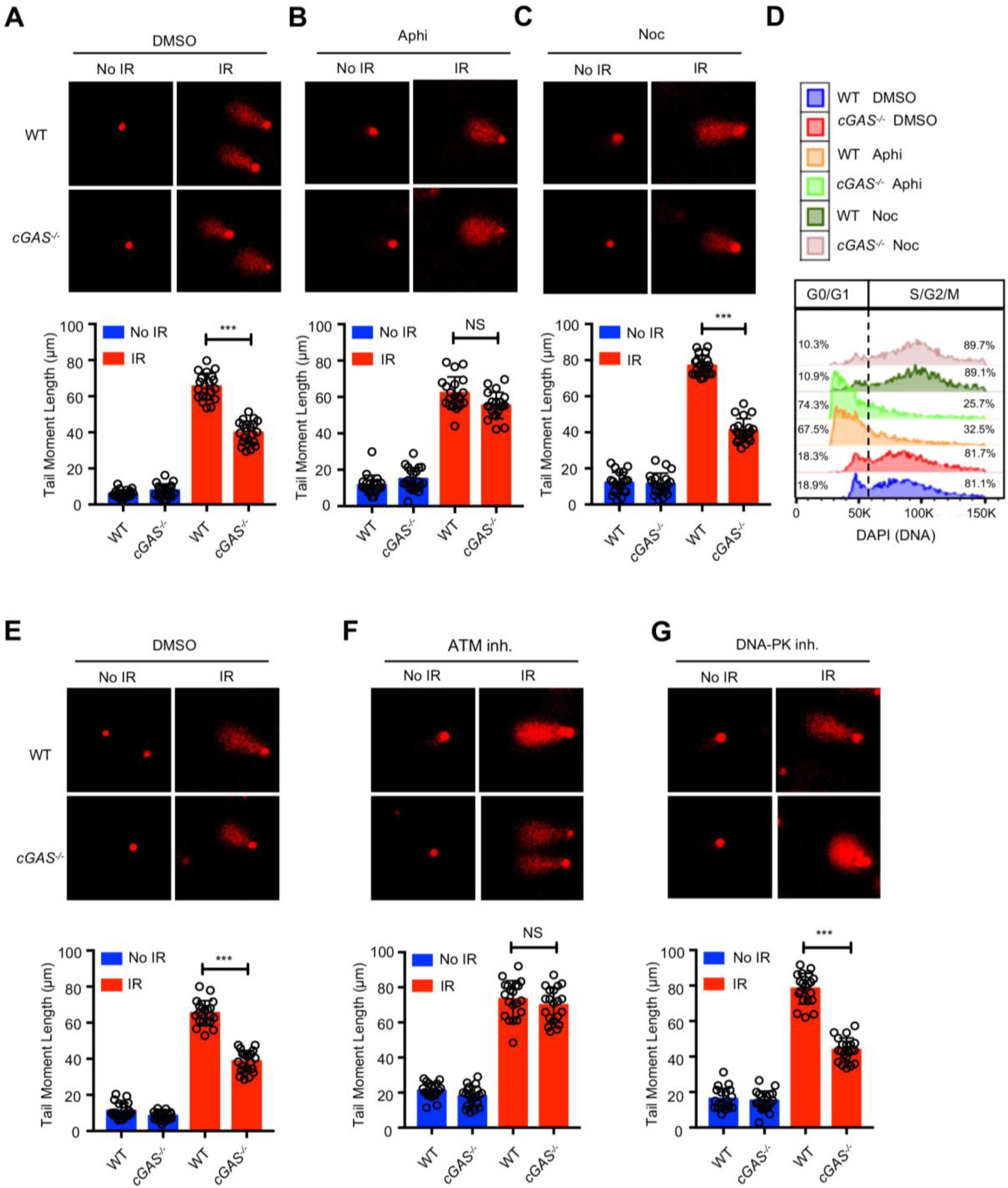
cGAS impedes DNA repair in a cell cycle and ATM-dependent manner. (**A-D**), cGAS suppresses DNA repair at G2 but not at G1/early S cell cycle phase. Representative images (upper) and corresponding quantification of comet tails in WT and *cGAS^−^/-* BMDMos that were either treated with (**A**) DMSO (control), (**B**), Aphidicolin (+Aphi) or (**C**) Nocodazole (+Noc) then γ-irradiated (IR: 10 Gy) and analyzed 15 min later. (**D**) Flow cytometric cell cycle analysis or cells in A-C. (**E-G**), Microscopic images (upper) and corresponding quantification of comet tails in WT and *cGAS^−^/-* BMDMos in (**A**) DMSO (control) or treated with (**F**) ATM inhibitor or (**G**) DNA-PKc inhibitor then γ-irradiated (IR: 10 Gy) and analyzed 15 min later. Each data set in the graphs represent mean score from 20 different fields with *n* > 200 comets; unpaired two-tailed Student’s t-test. NS: P >0.05, **P ≤ 0.01. For all figures, experiments were repeated at least three times in triplicates. Data are represented as Mean ± SEM.

### cGAS attenuates HR-DNA repair via DNA binding and oligomerisation

To independently verify the above findings, we examined the repair of a site-specific DSB induced by the I-SceI endonuclease using the direct repeat-GFP (DR-GFP) ^39^ and the total-NHEJ-GFP (EJ5-GFP) ^40^ reporter systems for HR and NHEJ, respectively. siRNA knockdown of endogenous cGAS in U2OS cells increased HR efficiency but minimally affected NHEJ repair (**Figure 5A-D**). In contrast, expression of hcGAS in HEK293T cells strongly reduced the efficiency of HR but not the NHEJ repair (**Figure 5E, F**).

**Figure 5.**
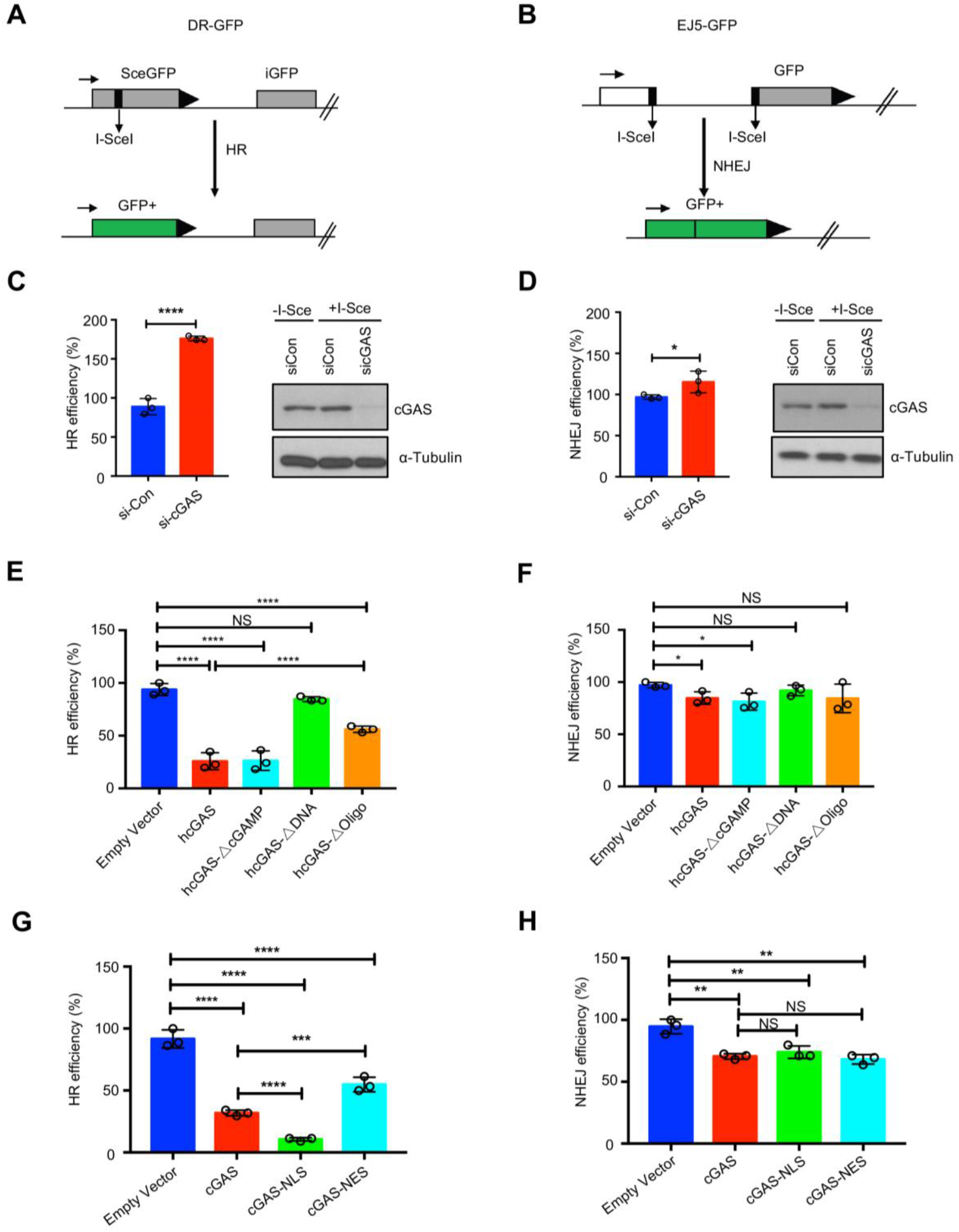
cGAS inhibits homologous recombination DNA (HR) via its DNA binding and oligomerization. (**A-B**), Schematics of HR and NHEJ reporter assays. (**C-D**) Obtained results showing the effect of silencing cGAS on HR (**C**) and NHEJ (**D**) in U2OS cells. Immunoblot inserts in C and D depict knockdown efficiency of cGAS. Tubulin: loading control. (**E-F**), Results showing the effect of hcGAS, hcGAS△cGAMP, or hcGAS△DNA or hcGAS△Oligo on HR (**E**) or (**F**) NHEJ in HEK293 cells. Data are means with S.E.M., n=3. Data are means with SEM., n=3, unpaired two-tailed Student’s t-test. **** P ≤ 0.0001. NS: P>0.05. (**G-H**), Reporter assays showing the effect of NLS and NES on cGAS-mediated inhibition of DNA repair.

To elucidate the cGAS features mediating inhibition of HR, we tested different cGAS mutants. The catalytically dead hcGASΔcGAMP inhibited HR to a similar degree as hcGAS. In contrast, the DNA binding (hcGASΔDNA) and oligomerization (hcGASΔOligo) mutants respectively, exhibited a complete or severe lack of such inhibitory effect (**Figure 5E, F**). These data conclusively demonstrate that cGAS specifically blocks HR-mediated DNA repair via its DNA binding and oligomerization but not catalytic activity. To further interrogate these findings, we addressed the importance of cGAS nuclear localization by testing the impact of hcGAS-NLS and hcGAS-NES on HR-DNA repair. Consistent with its increased accumulation in the nucleus (**Figure S2C**) cGAS-NLS had a stronger inhibitory effect on HR-DNA repair than hcGAS. In contrast, hcGAS-NES exhibited a weaker inhibitory effects (**Figure 5G, H**) consistent with its reduced presence in the nucleus (**Figure S2C, D**). Together, these results demonstrate that the ability of nuclear cGAS to impede HR-DNA repair depends on its DNA binding and oligomerization but not its enzymatic function.

### cGAS interferes with RAD51-mediated DNA strand invasion

To understand the specific signaling step in HR-DNA pathway targeted by cGAS we asked whether cGAS was inhibiting activation of the HR kinase ATM. However, expression of GFP-hcGAS or hcGASΔcGAMP in HEK293T or HEK293 cells had no effect on γ-irradiation-induced phosphorylation of ATM (**Figure S4B, C**). Similarly, WT, *cGAS^−/−^* and *Sting^−/−^* BMDMos showed comparable γ-irradiation-induced ATM activation (**Figure S4D**). To determine whether cGAS is recruited to DSB sites, we employed a DSB reporter system based on a mCherry-LacI-FokI nuclease fusion protein to create DSBs within a single genomic locus in U2OS cells (U2OS-DSB reporter) ^41^. GFP-hcGAS and mCherry-LacI-FokI did not co-localize at the DSB sites (**Figure S5A**). Moreover, GFP-hcGAS and γ-H2AX foci were not colocalized in γ-irradiated HEK293 cells (**Figure S5B**). Moreover, biochemical fractionation studies confirmed that cGAS was constantly in the nucleus and remained unchanged upon γ-irradiation (**Figure S5C**) indicating that observed phenotype was not via a cGAS effect on proximal signaling events at DSB sites but most likely due to inhibition of a critical downstream process.

The RAD51 recombinase acts downstream of ATM to catalyse HR-mediated DSB repair. Specifically, protomers of RAD51 form a protein filament on 3′ single-stranded DNA (ssDNA) tails stemming from the DSB end resection process ^37, 38, 42^. The RAD51-ssDNA filament, also referred to as the presynaptic filament, searches for and invades a homologous duplex target and exchanges ssDNA strands with the latter to generate a displacement loop (D-loop). This is followed by DNA synthesis and resolution of DNA intermediates to complete repair ^37, 38, 42^. Given the above data suggesting that cGAS inhibits a step downstream of ATM, we investigated whether cGAS might inhibit a RAD51-dependent process, first, by using the RAD51 Inhibitor B02, a small molecule that attenuates DNA binding by RAD51. As expected, treatment of cells with B02 caused an overall increase in fragmentation of genomic DNA.

Notably, the cGAS-mediated increase in fragmentation of genomic DNA was not detected in B02-treated in BMDMos and HEK293 cells (**Figure 6A, B and Figure S6A, B**), lending credence to the idea that cGAS might be interfering with a RAD51-controlled step in DSB repair. However, GFP-hcGAS was found not to colocalize with or impede the formation of RAD51 foci in γ-irradiated HEK293 cells (**Figure S6C**), again supporting the premise that cGAS was not inhibiting proximal signaling events at DSB sites.

**Figure 6.**
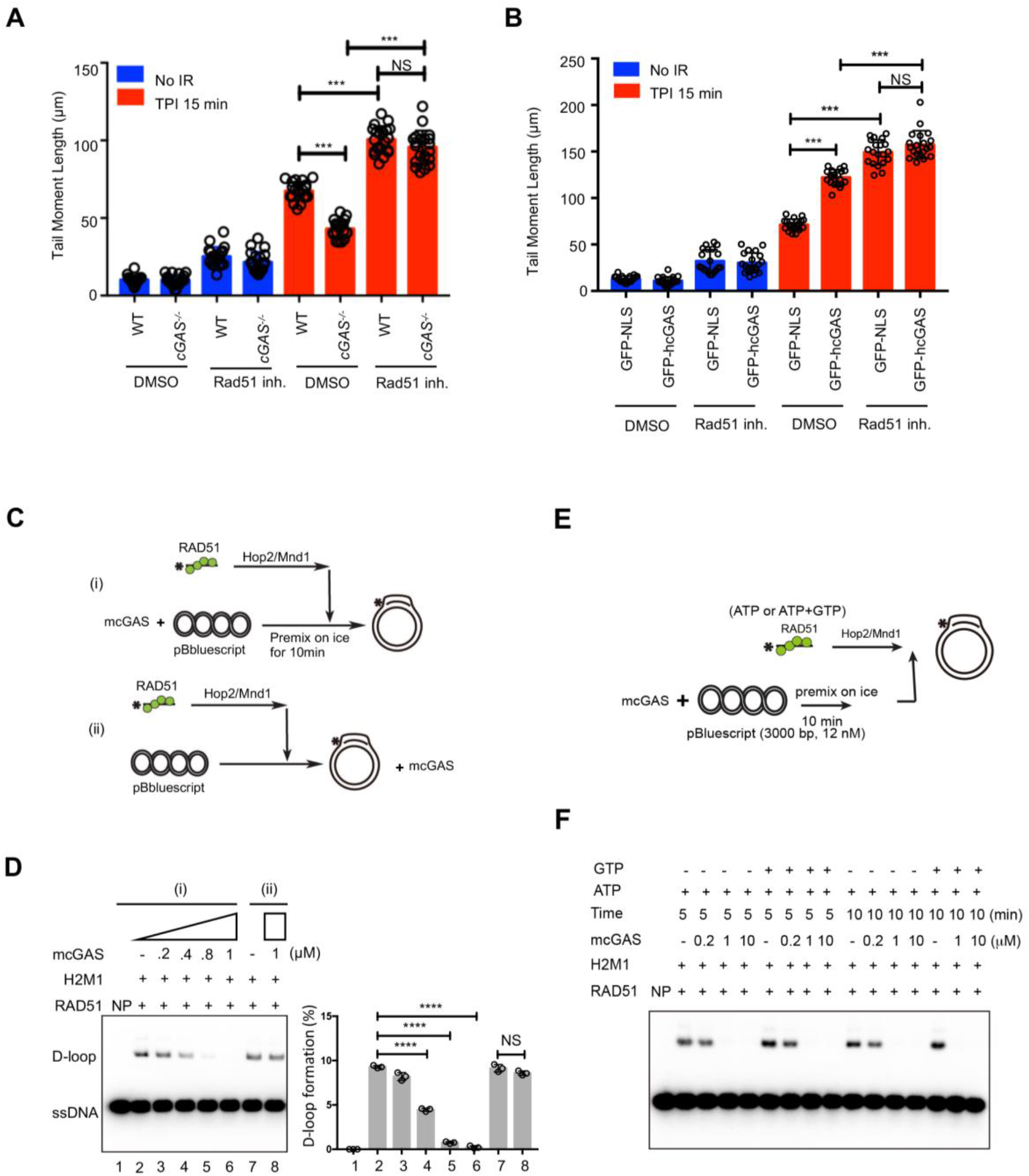
cGAS inhibits the HR DNA repair pathway by blocking RAD51-mediated D-loop formation. (**A-B**), RAD51 inhibition abrogates cGAS-mediated suppression of DNA repair. Comet tail moments of WT and *cGAS^−/−^* BMDMos(**A**) or GFP-NLS-, or GFP-hCGAS-expressing HEK293 cells (**B**) pretreated with the RAD51 inhibitor B02 (20 μM) 12 hour prior to γ-irradiation (10 Gy). Comet tail moments were quantified from 20 different fields with *n* > 200 comets; unpaired two-tailed Student’s t-test. NS: P >0.05, ***P ≤ 0.001. (**C**), Schematics of the D-loop formation assay, including pre-incubation of template dsDNA with cGAS (i), or with cGAS being added after RAD51 was bound to dsDNA (ii). (**D**) Pre-incubation of dsDNA with cGAS prevents D-loop formation by human RAD51, but does not affect the RAD1 activity once RAD51 filaments are bound to dsDNA. The percentage of D-loop formed in each reaction (left) was graphed as the average of triplicates ± SD., unpaired two-tailed Student’s t-test. NS P >0.05, ****P ≤ 0.0001. (**E**), Schematics of the D-loop assay. (**F**) Pre-incubation of template dsDNA with cGAS blocks subsequent D-loop formation regardless of the presence of cGAMP precursors (ATP+GTP).

Condensation of DNA into higher-ordered structures is a barrier to HR ^43, 44^. cGAS induces the formation of higher-ordered ladder structures via its DNA binding and subsequent oligomerization ^45-47^. In view of this and the above data showing that inhibition of HR by cGAS was dependent on its DNA binding and oligomerization, we tested whether cGAS was hindering RAD51 filaments from invading target homologous dsDNA template. To test this premise, we examined the effect of purified mouse cGAS (**Figure S7A**) in the RAD51-mediated D-loop reaction with ssDNA and supercoiled dsDNA as substrates (schematics, **Figure 6C, E**). Incubation of the supercoiled DNA template with cGAS led to a strong inhibition of D-loop formation (**Figure 6D** compare lane 2 vs 4-6). However, if RAD51 filaments were pre-bound to template dsDNA (schematic **Figure 6C (ii)**) then cGAS did not interfere with RAD51-mediated D-loop formation (**Figure 6D**, compare lane 7 vs 8). This demonstrates that cGAS-mediated attenuation of D-loop formation was not due to inhibition of the enzymatic activity of RAD51 but by hindering the RAD51 filaments from accessing the dsDNA template. Accordingly, when pre-incubated with linear dsDNA template, cGAS also inhibited RAD51-mediated DNA strand exchange between ssDNA and the linear dsDNA (**Figure S7B-D**). Noteworthy, in both assay systems, inhibition occurred regardless of whether ATP and GTP, the precursors required for cGAS-mediated cGAMP synthesis, were present in the reaction, thus confirming that the inhibitory activity of cGAS is not cGAMP-mediated (**Figure 6E, F**). Of note, cGAS also inhibited DNA strand exchange mediated by yeast Rad51 protein (**Figure S7E, F**).

Finally, we asked whether inhibition of RAD51-mediated D-loop formation is a universal feature of dsDNA binding proteins. First, we tested for a protein that binds dsDNA with a similar affinity as cGAS and identified MHF, a component of the Fanconi anemia (FA) core complex ^48^ as protein fitting this criteria (**Figure S8A, B**). Remarkably, in spite of comparable binding to dsDNA, in sharp contrast to cGAS, we found that MHF does not inhibit D-loop formation by human RAD51 (**Figure S8C-E**), thus demonstrating that observed inhibition of D-loop formation is not non-specific but an inherent feature of cGAS owed to both its DNA binding and oligomerization.

All together these results demonstrate that cGAS is a negative regulator of HR-mediated DSB repair, and we have provided biochemical evidence that cGAS acts by inhibiting RAD51-mediated DNA strand invasion.

## DISCUSSION

Whereas accurate repair of DNA lesions via HR is indispensable for organismal integrity, unrestrained HR may lead to undesired endpoints including chromosomal translocation, deletion, inversion or loss of heterozygosity. Therefore, to match specific cellular needs, HR is subject to careful regulation via mechanisms not fully understood. Since its discovery a few years ago ^17^, cGAS has been described as a cytoplasmic sensor that alerts the innate immune system to the presence of foreign or self DNA, leading to not only antimicrobial and antitumor immunity, but also autoimmunity, neurodegeneration and cancer ^10, 28^. Here we have uncovered a new function for cGAS in the negative regulation of HR, and that this property contributes to genome destabilization. We show that this nuclear property of cGAS is uncoupled from its enzymatic activity or the STING-IFN-I pathway. A model incorporating this new function and the known cytoplasmic role of cGAS in innate immunity is shown in **Figure 7**.

**Figure 7.**
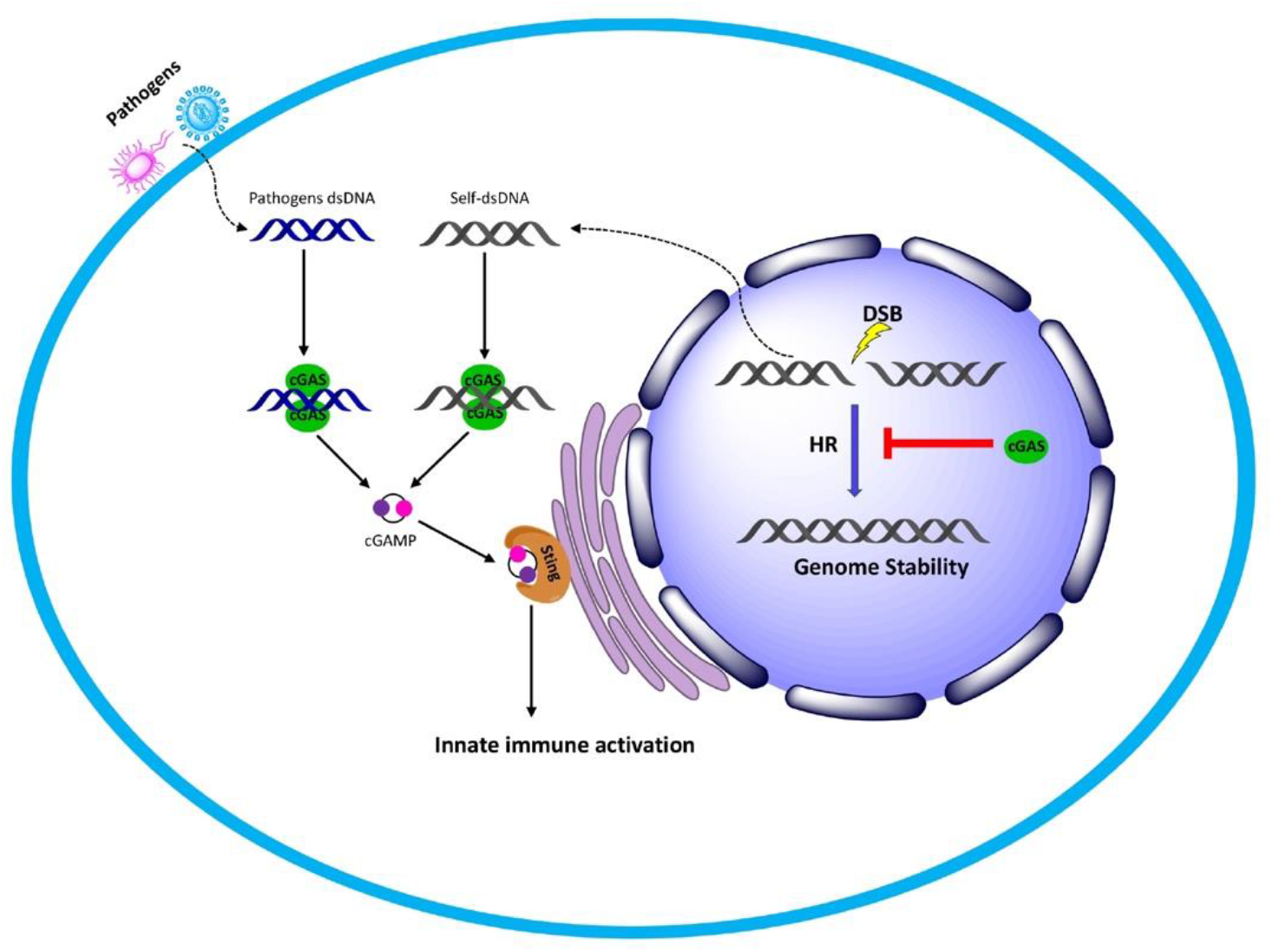
Model of the distinct subcellular functions of cGAS. Aside from the canonical function in the cytoplasm in sensing microbial- or self-DNA to activate the innate immune system, our work reveals that in the nucleus cGAS inhibits homologous DNA repair thereby promotes genome destabilization. This later function requires the DNA binding of cGAS but not its enzymatic activity and the downstream STING signaling events.

Mechanistically, we show that, cGAS hinders RAD51-mediated strand invasion, a critical step in HR. This mechanism of regulation is perhaps analogous to that by DNA compaction proteins such as the linker histone H1^43, 44, 49^ proposed to inhibit HR by limiting access of RAD51 filaments to template DNA ^50^. What is the biological relevance of cGAS-mediated attenuation of HR? We posit that under homeostatic conditions, cGAS may generally function as a negative regulator to suppress unscheduled genome rearrangement to prevent chromosomal aberrations. On the other hand, by inhibiting HR-DNA repair in cells under severe genotoxic stress, cGAS promotes mitotic catastrophic death thereby limiting the transfer of defective genomes to the next generation.

Defects in DNA repair and immune pathways are relevant for many human afflictions including infections, autoimmunity, neurodegeneration, cancer and aging-associated disorders. Recently, there is a growing appreciation that the DNA damage response and the immune system are interconnected, and that a key molecule at the intersections of these biological systems is cGAS. All the biological effects linked to cGAS has been ascribed to its role as an innate immune sensor of DNA. We have now established that independent of its canonical function in immune activation, cGAS is a negative the regulator of DNA repair hence accelerates the cellular outcomes of DNA damage.

This study presents a new entry point towards a better understanding of how cGAS may impact upon a variety of diseases, in particular those associated with genome instability and immune dysfunctions. For example, cGAS is increasingly acknowledged to promote both tumorigenesis ^11, 51, 52^ and anti-tumor immunity ^4-6, 53^. Understanding the extent and biological context in which these distinct subcellular functions of cGAS contribute to anti-tumor immunity or drive tumorigenesis, and how they can be manipulated will be beneficial for achieving the desired outcome of DNA damage- and immune-based anti-tumor therapies.

## MATERIALS AND EXPERIMENTAL PROCEDURES

### Mice

All the mice in this study were on pure C57BL/6 background. *Sting^−/−^* (C57BL/6J-Tmem173gt/J) ^54^ and *cGAS^−/−^* (B6(C)-Mb21d1tm1d (EUCOMM) Hmgu/J) ^55^ were from Jackson Laboratory. Mice were bred in specific pathogen-free animal facility of Umeå center for comparative Biology (UCCB) and experiments carried out according to the guidelines set out by the Umeå Regional Animal Ethic Committee (Umeå Regionala Djurförsöksetiska Nämnd), Approval no. A53-14.

### Antibodies and Reagents

The anti-α-Tubulin antibody, aphidicolin, nocodazole, the ATM inhibitor (KU-55933), DNA-PK inhibitor (NU7026) and the Rad51 inhibitor (B02) were purchased from Sigma-Aldrich. The anti p-ATM (Ser1981) and GFP antibody was from Santa Cruz. Antibodies against ATM, Flag, mouse cGAS, human cGAS, STING, H2A, H2A.X, γ-H2A.X p-IRF3 and IRF3 were from Cell Signaling Technology. 2’,3’-cGAMP and Immunostimulatory DNA (ISD) were from InvivoGen. ATP was from New England Biology while GTP, Rad51 and Lamin B1 antibody was from Abcam.

### Plasmids and constructs cloning

pTRIP-SFFV-EGFP-NLS(GFP-NLS), pTRIP-CMV-GFP-FLAG-hcGAS(GFP-hcGAS) and pTRIP-CMV-GFP-FLAG-hcGAS E225A-D227A(GFP-hcGAS(ΔcGAMP) (Addgene plasmid #86677, #86675,and #86674) have been described previously ^30^. pTRIP-CMV-GFP-FLAG-hcGAS C396A-C397A(GFP-hcGAS(ΔDNA)) and pTRIP-CMV-GFP-FLAG-hcGAS K394E were generated by site directed mutagenesis from pTRIP-CMV-GFP-FLAG-hcGAS(GFP-hcGAS). Flag-hcGAS was cloned into pcDNA3.1+ to generate the pcDNA-hcGAS plasmid. pcDNA-hcGAS E225A-D227A(pcDNA-hcGAS(ΔcGAMP), pcDNA-hcGAS C396AC397A(GFP-hcGAS(ΔDNA) and pcDNA-hcGAS E394A were generated by site directed mutagenesis from the pcDNA-hcGAS plasmid. The SV40 NLS (nuclear localization signal) sequence (5’-CCAAAAAAGAAGAGAAAGGTA-3’) was cloned separately into C terminal of pTRIP-CMV-GFP-FLAG-hcGAS and pcDNA-hcGAS to generate pTRIP-CMV-GFP-FLAG-hcGAS-NLS and pcDNA-hcGAS-NLS. NES (nuclear export signal) sequence (5’-CTGCCCCCCCTGGAGCGCCTGACCCTG-3’) was cloned separately into C terminal of pTRIP-CMV-GFP-FLAG-hcGAS and pcDNA-hcGAS to generate pTRIP-CMV-GFP-FLAG-hcGAS-NES and pcDNA-hcGAS-NES. pHPRT-DRGFP and pCBASceI were gifts from Maria Jasin (Addgene plasmid # 26476 and # 26477) ^39^. pimEJ5GFP was a gift from Jeremy Stark (Addgene plasmid # 44026) ^40^.

### Cells and cell culture

HEK293, HEK293T, U2OS cells were cultured under 5% CO2 at 37°C in Dulbecco’s modified Eagle medium (DMEM, high glucose, GlutaMAX) (Life Technologies) containing 10% (v/v) fetal calf serum (FCS, GIBCO), 1% (v/v) penicillin (100 IU/ml)+streptomycin (100 μg/ml). Bone-marrow-differentiating monocytes (BMDMos) were generated by culturing the mouse bone marrow cells in IMDM medium (GIBCO, Life Technologies) supplemented with 10% (v/v) FCS (GIBCO, Life Technologies), 1% (v/v) penicillin (100 IU ml−1)/streptomycin (100 μg/ml), 2 mM glutamine (Sigma-Aldrich) and 20% (v/v) L929 conditional medium and maintained with 5% CO2 at 37 °C. The cells were used for experiment on 4 days after start of differentiation.

### Generation of stable cell line

HEK 293T cells were transfected with psPAX2, pMD2.G plasmids and the lentiviral vector pTRIP containing the open reading frame of GFP-NLS or GFP-cGAS or GFP-cGAS mutants by using Lipofectamine LTX. The supernatants containing lentiviral particles were harvested at 48 h. HEK293 and HEK293T cells were then transduced with the lentiviral vectors by directly adding supernatant together with polybrene (5ug/ml) to cells. 2 days later, GFP positive cells were sorted by flow cytometry and propagated further. To generate stable HEK293T DNA damage reporters HEK293T cells were transfected separately with pHPRT-DRGFP (to monitor HR) and pimEJ5GFP (to monitor NHEJ), 2 days later cells were put under puromycin (2ug/ml) selection. Single clones were picked and expanded for the reporter assays.

### Immunofluorescence

Cells were seeded and cultured on glass coverslips in 12 well plate and fixed in 4% paraformaldehyde (PFA) in PBS for 20 min at room temperature. Cells were permeabilized in 0.5% Triton X-100 for 10 min. Slides were blocked in 5% normal goat serum (NGS), and incubated with primary antibodies diluted in 1% NGS overnight at 4°C. Samples were then incubated with secondary antibodies labeled with Alexa Fluor 488 (Invitrogen) diluted in 1% NGS at RT for 1 h.

Thereafter they were stained with DAPI (or plus Phalloidin) for 15 min at room temperature. Coverslips were mounted using Dako fluorescence mounting medium (Agilent) and imaged using Nikon confocal (Eclipse C1 plus). All scoring was performed under blinded conditions.

### Subcellular fractionation and immunoblotting

For cytoplasmic and nuclear extracts were prepared using the nuclear extraction kit (Abcam) according to the manufacturer’s instructions. For chromatin bound fraction, we use the Subcellular Protein Fractionation Kit (Thermo Fisher) according to the manufacturer’s instructions. For other assays, cells grown in culture were trypsinized, pelleted, washed, and resuspended in a mild Nonidet P-40 lysis buffer (1% NP-40, 50 mM Tris-HCl, 150 mM NaCl pH 7.5, 1 mM NaF, 2 mM PMSF, protease inhibitor cocktail [Roche Applied Science], 1 mM sodium orthovanadate and 10 mM sodium pyrophosphate). The lysates were centrifuged at 13,000 rpm for 15 min, and proteins in supernatants were quantified by BCA reagent (Thermo Fisher Scientific, Rockford, IL). Proteins were resolved in SDS-PAGE, transferred to nitrocellulose membrane (Amersham protan 0.45µm NC) and immunoblotted with specific primary antibodies followed by HRP-conjugated secondary antibodies. Protein bands were detected by Supersignal West Pico or Femto Chemiluminescence kit (Thermo Fisher Scientific).

### Mitotic catastrophic death

BMDMos were arrested at G2 by incubation with 100 nM nocodazole for 12 hours. Thereafter they were γ-irradiated then released into mitosis and evaluated for cell death at indicated time points. Irradiation-induced mitotic catastrophic death was determined by XTT assay (Sigma-Aldrich) according to the manufacturer’s instructions. Absorbency was measured with a spectrophotometer (Tecan Infinite M200 Microplate reader) at 450 nm with a reference wavelength at 650 nm. Relative number of dead cells as compared to the number of cells without treatment was expressed as percent mitotic cell death using the following formula: mitotic cell death (%)=100% - 100%(A450 of treated cells/A450 of untreated cells).

### Comet assay

Cells were γ-irradiated in a 137Cs gamma-ray source (Gammacell 40 irradiator, MDS Nordion) with indicated dose and chromosome fragmentation was determined by comet assay. Briefly, during irradiation all the cells are keep in ice to stop the DNA repair process. Thereafter, cells were transfer to 37℃ to allow DNA repair and then harvested at indicated time points for analysis. 1 × 10^5^ cells/ml in cold PBS were resuspended in 1% low-melting agarose at 40°C at a ratio of 1:3 vol/vol and pipetted onto a CometSlide. Slides were then immersed in prechilled lysis buffer (1.2 M NaCl, 100 mM EDTA, 0.1% sodium lauryl sarcosinate, 0.26M NaOH PH>13) for overnight (18-20 h) lysis at 4°C in the dark. Slides were then carefully removed and submerged in room temperature rinse buffer (0.03M NaOH and 2mM EDTA, pH > 12) for 20 min in the dark. This washing step was done 2 times.

Slides were transferred to a horizontal electrophoresis chamber containing rinse buffer and separated for 25 min at voltage (0.6V/cm). Finally, slides were washed with distilled water and stained with 10 µg/ml propidium iodide and analyzed by fluorescence microscopy. 20 fields with about 200 cells in each sample were evaluated and quantified by the Fiji software to determine the tail length (tail moment).

### Pulsed-field gel electrophoresis

BMDMos from WT and *cGAS^−/−^* mice were irradiated (20 Gy or 30 Gy) on ice (time 0) then incubated at 37°C to allow DNA repair. At different time points post irradiation (15, 45 min) cells were washed twice with ice-cold phosphate-buffered saline (PBS). Cell pellets (plugs) were immediately placed in 10x volume of lysis buffer (0. 5MEDTA (pH 9.5), 1 % sarkosyl, and 1 mg/ml proteinase K (Sigma) for a 48-h digestion at 50°C with one buffer change after 24 h. Following lysis, the plugs were washed for at least 24 h with 10x volume of TE buffer containing 10mM phenylmethylsulphonyl fluoride (PMSF) at room temperature. After PMSF treatment, the plugs were washed three times for at least 2 h in each case in 10x volume of TE buffer without PMSF at room temperature.

Electrophoresis was performed in a CHEF-DR II apparatus (Bio-Rad) with a hexagonal array of 24 electrodes, which produce a field reorientation angle of 120°. The plugs were inserted into 1% gels made from high tensile strength agarose (pulsed field grade agarose; Bio-Rad) in 0.5x TBE. The gel was run at 13°C in 0.5x TBE (the buffer was recirculated through a refrigeration unit to keep the temperature constant and to avoid ion build up at the electrodes) for 36 h. The pulse time was increased during the run linearly from 50 to 150 s at a field strength of 6 V/cm. After electrophoresis, the gels were stained for 1 h in 200 ml staining buffer, (TE containing 10 µg/ml ethidium bromide) and de-stained for 3 h in the same buffer in the absence of ethidium bromide. After that, the signals were detected by a Gel-DOC system.

### Determination of micronuclei

HEK293 cells exposed (or not) to γ-irradiation and cultured for 24 hours. BMDMos arrested at G2 by incubating with Aphidicolin were to γ-irradiation then cultured for 48 hours. Cells were fixed, permeabilized in 0.5% Triton X-100, stained with the DNA dye DAPI, then analysed by microscopy for the presence of micronuclei. Micronuclei were defined as discrete DNA aggregates separate from the primary nucleus in cells where interphase primary nuclear morphology was normal. Cells with an apoptotic or necrotic appearance were excluded.

### HR and NHEJ reporter assays

Homologous recombination (HR) and NHEJ repair in HEK293T cells was measured as described previously using the DR-GFP stable cells ^39^ and EJ5-GFP stable cells ^40^. Briefly, 0.5 × 10^6^ HEK293T stable reporter cells were seeded in 6-well plates. co-transfected with 2 μg I-SceI expression plasmid (pCBASce) and either 4 μg pcDNA-hcGAS, pcDNA-hcGASΔcGAMP pcDNA-hcGAS-ΔDNA or empty pcDNA vector. 48 hours post transfection, cells were harvested and analysed by flow cytometry analysis for GFP expression. Means were obtained from three independent experiments. U2OS cells silenced for cGAS using were transfected with 2 μg I-SceI expression plasmid (pCBASce) for 2 days then harvested and analysed by flow cytometry analysis for GFP expression.

### Protein purification

Purified mouse cGAS (mcGAS aa 141–507) was a gift from Karl-Peter Hopfner, and the method for purification has been described ^47^. Human RAD51, Hop2/Mnd1, MHF and budding yeast Rad51 and Rad54 were purified as described previously ^48, 56-58^.

### D-loop formation

The D-loop reaction was conducted as described previously ^59^. Briefly, mouse cGAS protein (0.2-1.0 μM) was pre-incubated with pBluescript dsDNA (36 μM base pairs) on ice for 10 min. Human RAD51 (0.6 μM) was incubated with 32P-labeled 90-mer ssDNA (2.4 μM nucleotides) at 37°C for 10 min to allow RAD51 filament formation. Hop2/Mnd1 complex (0.5 μM) was then added to the mixture, followed by a 2-min incubated at 37°C. The reaction was initiated by adding the cGAS-pBluescript dsDNA mixture and further incubated at 37°C for 5 min. The reaction mixtures were deproteinized before being resolved in 0.9% agarose gels in TBE buffer. Gels were dried and the radiolabeled DNA species were revealed and quantified by phosphorimaging analysis. cGAS protein was also added after D-loop formation, followed by a further 5-min incubation at 37°C.

### DNA strand exchange assay

The assay was conducted at 37°C and reaction mixtures were resolved by electrophoresis in non-denaturing 10% polyacrylamide gels in TAE buffer (45 mM Tris-acetate, pH 7.5, 0.5 mM EDTA) as described previously ^60^. Briefly, the 150-mer oligo (6 μM nucleotides, P1 in Table S1) was incubated with human RAD51 (2 μM) in 10 μl of buffer G (25 mM Tris-HCl, pH 7.5, 60 mM KCl, 1 mM DTT, 100 μg/ml BSA, 1 mM ATP/1 mM GTP, and 2 mM MgCl2) containing an ATP-regenerating system consisting of 20 mM creatine phosphate and 20 μg/ml creatine kinase for 5 min. cGAS was premixed with 32P-labeled homologous dsDNA (6 μM base pairs, P2/P3 in Table S1) on ice for 10 min. The two reaction mixtures were combined, followed by the addition of 4 mM spermidine hydrochloride to 12.5 μl final volume. After 30 min of incubation, the reactions were stopped by adding an equal volume of 1% SDS containing proteinase K (1 mg/ml) and a 5-min incubation. Gels in which the deproteinized reaction mixtures had been resolved were dried and subject to phosphorimaging analysis.

### DNA mobility shift assay

cGAS (20 to 200 nM) or MHF hetero-tetramer ^48^ (20 to 200 nM) was incubated with an 80 mer double strand DNA substrate (dsDNA, 10 nM each) at 37°C for 10 min in 10 μl of buffer B (25 mM Tris-HCl, pH 7.5, 1 mM DTT, 100 μg/ml BSA, 1 mM MgCl_2_, and 45 mM KCl). The reaction mixtures were resolved in 7% polyacrylamide gels in TAE buffer (40mM Tris, 20mM Acetate and 1mM EDTA) at 4°C. Gels were dried onto Whatman DE81 paper (Whatman International Limited) and subject to phosphorimaging analysis.

### RT-qPCR

Total RNA was extracted using the Trizol (Thermo Fisher) according to the manufacturer’s protocol. cDNA was prepared using Maxima H Minus First Strand cDNA Synthesis Kit and random oligomer primers (Thermo Fisher Scientific). qRT–PCR was performed using SYBR Select Master Mix (Thermo Fisher Scientific) on an QuantStudio 5 Real-Time PCR System (Thermo Fisher). The *IFNB1* transcript levels were normalized to the housekeeping gene 18S rRNA.

**Table S1.**
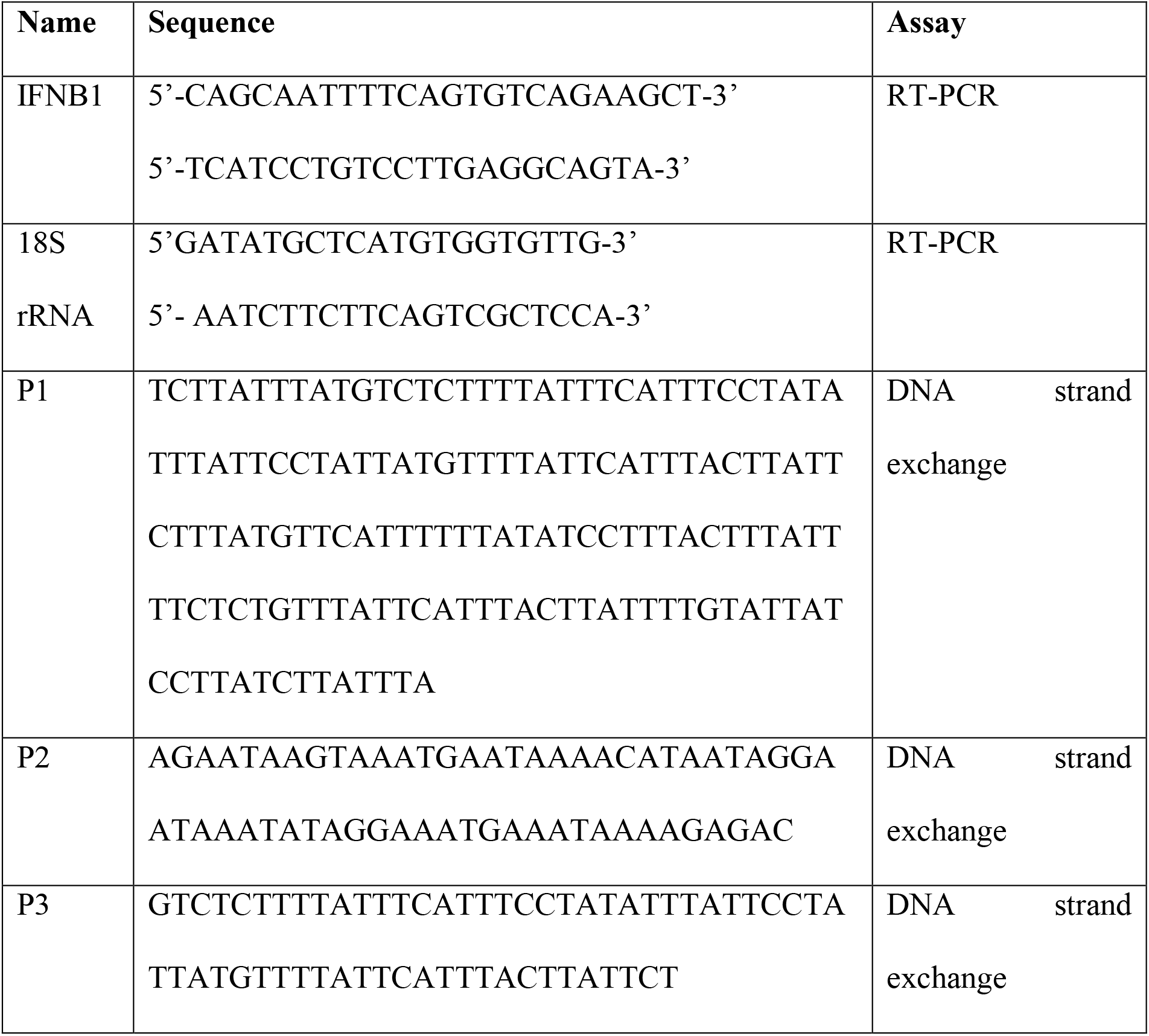
Oligonucleotides used in this study.

### siRNA-mediated cGAS silencing

To silence cGAS, U2OS cells were transfected with a pool of the following siRNA from Thermofischer (sicGAS-1: 5′-GGAAGAAAUUAACGACAUU-3′;sicGAS-2: 5′-GAAGA AACAUGGCGGCUAU-3′).

### FokI induced double strand breaks system

The U2OS-FokI DSB reporter cells contain a stably integrated LacO array and an mCherry-LacI-FokI fusion protein fused to a destabilization domain (DD) and a modified estradiol receptor (ER) (ER-mCherry-LacI-FokI-DD)^41^. This enables inducible nuclear expression of ER-mCherry-LacR-FokI-DD after administration of the small molecule Shield-1 ligand (stabilizes the DD-domain) and 4-hydroxytamoxifen (4-OHT; induce nuclear translocation of ER-mCherry-LacR-FokI-DD). To induce site-specific double-strand breaks by FokI, these cells were incubated with 1 µM Shield-1 (cat. no. 632189, Clontech) and 1 µM 4-OHT (cat. no. H7904, Sigma-Aldrich) for about 5h.

## Author contributions

H.J and N.O.G conceived the study. H.J., S. P., X.X., F.L., P.S., N.O.G, designed experiments and interpreted data. H.J., S. P., X.X, F.L. performed experiments. N.O.G supervised the research and together with H.J. wrote the paper which other authors commented on.

## Acknowledgments

We are grateful to Karl-Peter Hopfner, Ludwig-Maximilians-University, Munich, for providing recombinant murine cGAS. **Funding:** This work was funded by the Laboratory for Molecular Infection Medicine Sweden (MIMS), the Medical Faculty, Umeå University, the Swedish Research Council (grants 2015-02857 and 2016-00890 to N.O.G), the Swedish Cancer Foundation (grant, CAN 2017/421 to N.O.G) and the National Institutes of Health (NIH) (grant RO1 CA220123 to P.S.).

## Supplemental Information

**Figure S1.**
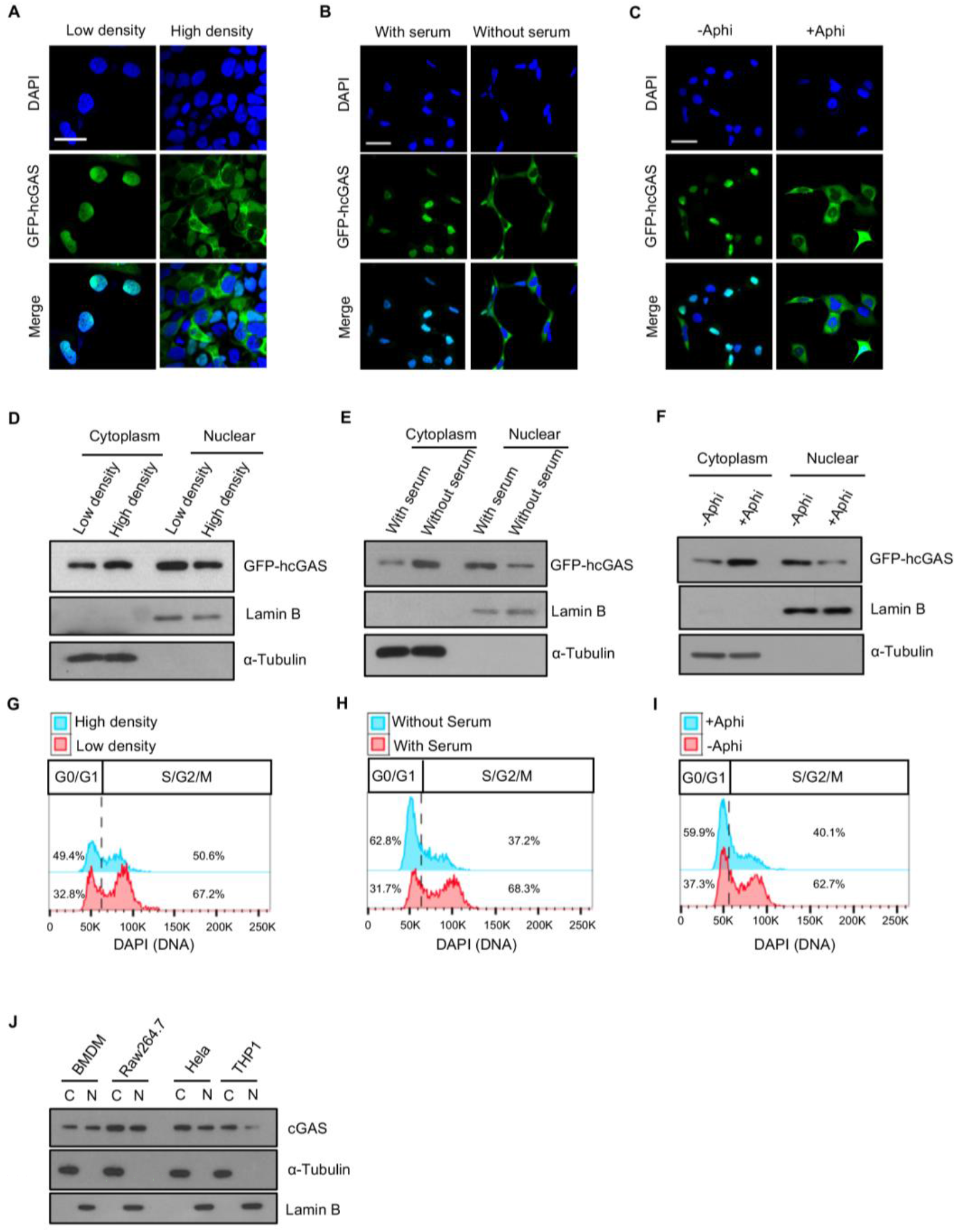
cGAS is constantly present in the cytosol and nucleus and is impacted by cell cycle. **(A-C)**, Fluorescence images of GFP-hcGAS in HEK293 cells cultured at (**A**) low/high density, (**B**) with/without serum, (**C**) with/without Aphidicolin (Aphi). DAPI (DNA). Scale bar: 100 μm. (**D-F)**, Immunoblots estimation of GFP-hcGAS in nuclear/cytosolic fractions of HEK293 cells cultured as in A-C. Lamin B and α-Tubulin are nuclear and cytosolic markers respectively. (**G-I**), Flow cytometric analysis of cell cycle of HEK293 cells depicted in DF. (**J**), Estimation of endogenous cGAS in nuclear/cytosolic fractions of Bone marrow derived macrophages (BMDMs), RAW 264.7 macrophages, HeLa cells and THP1 monocytes. Related to Figure1.

**Figure S2.**
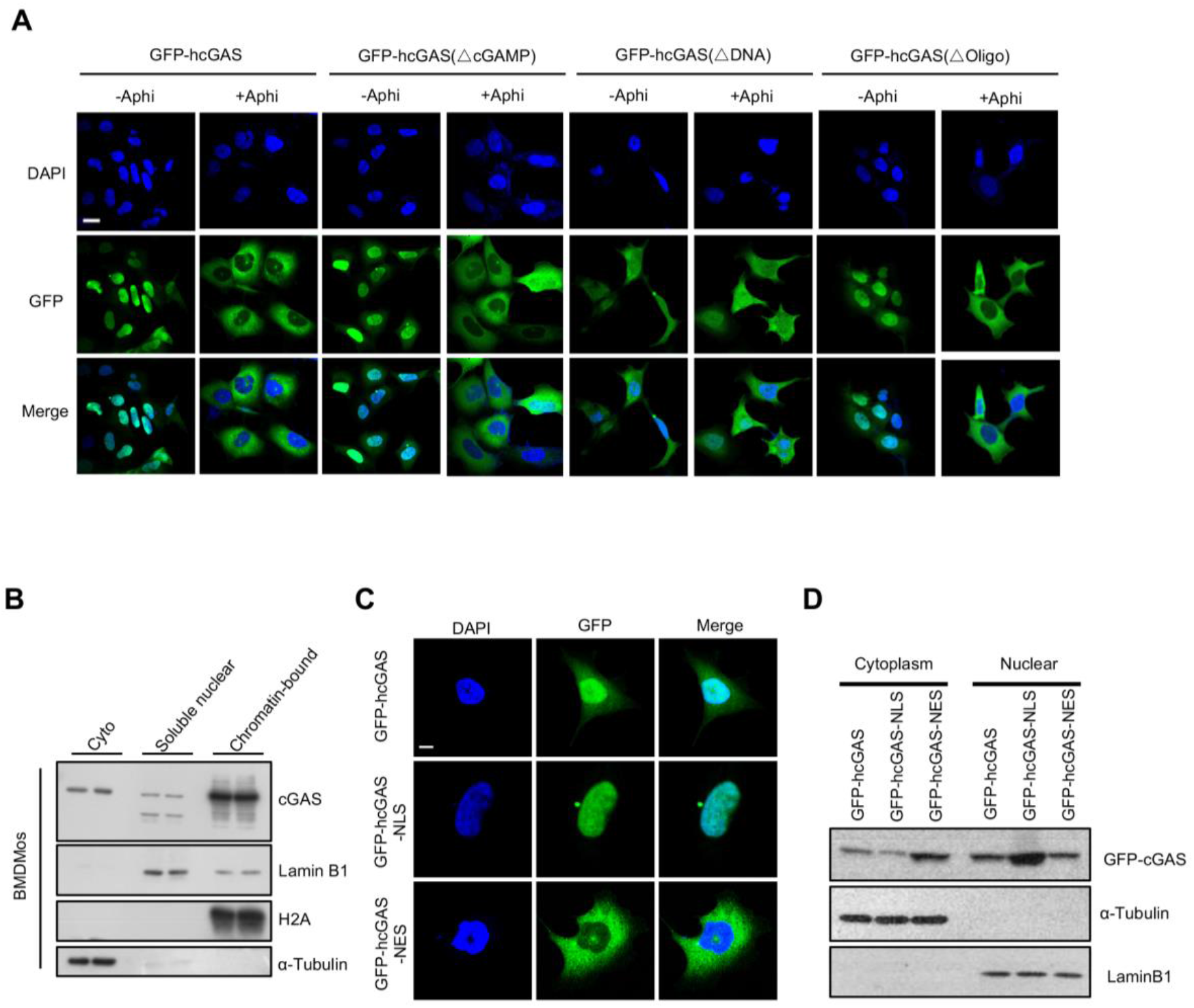
Localization and retention of cGAS in the nucleus is due to is avid binding to DNA. (**A**) Fluorescence images of GFP-hcGAS, GFP-hcGASΔcGAMP, GFP-hcGASΔDNA and GFP-hcGASΔOligo in HEK293 cells cultured with or without Aphidicolin. (**B**) Most of the nuclear cGAS is bound to chromatin: Cytosolic (cyto), soluble nuclear and chromatin fractions from BMDMos were immunoblotted for cGAS, (**C-D**) A nuclear export signaling (NES) is not sufficient to dislodge chromatin-bound cGAS from the nucleus. (**C**) Fluorescence images of GFP-hCGAS, GFP-hCGAS-NLS and GFP-hCGAS-NES in HEK293 cells. (**D**) Immunoblots of subcellular fractions of GFP-hCGAS-, GFP-hCGAS-NLS- and GFP-hCGAS-NES-expressing HEK293 cells.

**Figure S3.**
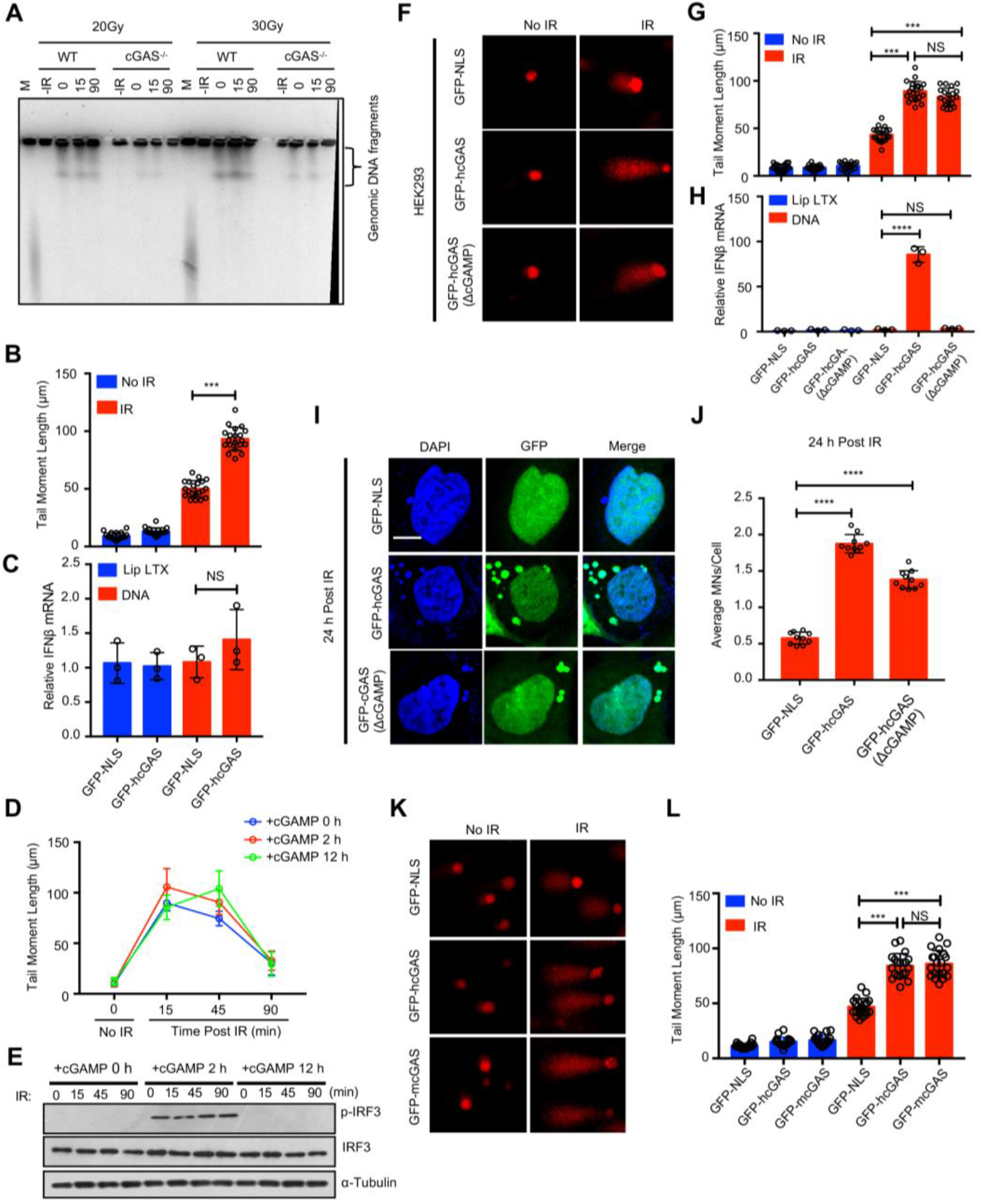
STING signaling is dispensable for cGAS-mediated inhibition of DNA repair but boosts micronuclei generation. (**A**), Pulsed-field gel electrophoresis analysis of WT and *cGAS^−^/-* BMDMos γ-irradiated (10 Gy) on ice then incubated at 37°C to allow DNA repair for indicated duration. (**B**), Comet tail moments in GFP-NLS- and GFP-hcGAS-expressing HEK293T cells γ-irradiated (IR: 10 Gy) on ice then incubated at incubated at 37°C for indicated duration. (**C**), RT-PCR analysis of *IFNB1* response in GFP-NLS- or GFP-hcGAS-expressing HEK293T cells stimulated with transfected DNA for 6 hours. (**D**), Comet tail moments of HEK293 cells stimulated with 10 ug/ml cGAMP for indicate periods then γ-irradiated and incubated at 37°C for indicated duration. (**E**), Immunoblots of IRF3 phosphorylation in HEK293 cells treated as in (D). (**F-G**), Microscopic images (**F**) and quantifications (**G**) of comet tails 15 min after irradiation of GFP-NLS-, GFP-hcGAS-, GFP-hcGASΔcGAMP) expressing HEK293 cells. (**H**), RT-PCR analysis of *IFNB1* response in GFP-NLS- or GFP-hcGAS-expressing HEK293 cells stimulated with transfected DNA for 6 hours. (**I**), Micronuclei in GFP-NLS-, GFP-hcGAS-, GFP-hcGAS(ΔcGAMP)- expressing HEK293 cells 24 hours after γ-irradiation (IR; 10 Gy). DAPI (DNA). Scale bar:10 μm. (**J**), Correspondingq uantification of micronuclei (MN)/cell. (**K-L**), Microscopic images (**F**) and quantifications of comet tails (**K**) 15 min after irradiation (10Gy) of GFP-NLS-, GFP-hcGAS-, or GFP-mcGAS-expressing HEK293 cells. Each data set bar comet graph was calculated from six different microscopic fields with over 200 cells; unpaired two-tailed Student’s t-test. **** P ≤ 0.0001. Mean ± SEM. of n=3 independent experiments; unpaired two-tailed Student’s t-test. NS P >0.05.

**Figure S4.**
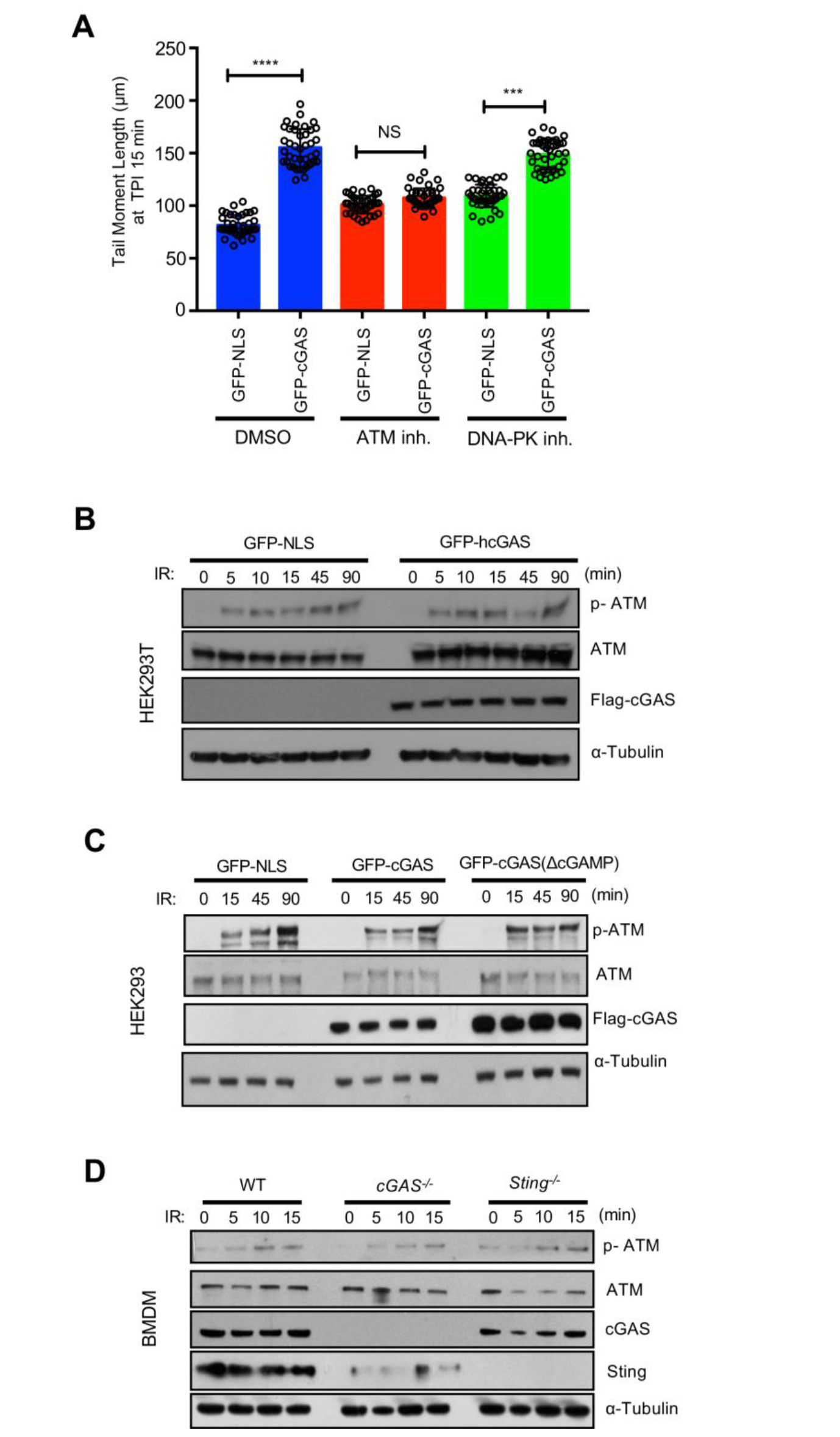
cGAS suppresses DNA repair in an ATM-dependent manner but not ATM activation. (**A**) Inhibition of the ATM kinase but not DNA-PKc abrogates cGAS-mediated suppression of DNA repair. Comet tail moments in GFP-NLS-, or GFP-hCGAS-expressing HEK293 cells pretreated for 12 hours with ATM inhibitor KU55933 (ATM inh.), 200 nM DNA-PK inhibitor NU7026 (DNA-PK inh.) then γ-irradiated (IR: 10Gy) and analyzed 15 min later. Data set represent mean score from 40 different fields with *n* > 200 comets; unpaired two-tailed Student’s t-test. NS: P >0.05, ***P ≤ 0.001. (**B-D**), cGAS does not impeded ATM activation. ATM phosphorylation in γ-irradiated (10 Gy) (**B**), GFP-NLS-, GFP-hcGAS-expressing HEK293T cells, or (**C**), GFPNLS-, GFP-hcGAS-, GFP-hcGASΔcGAMP-expressing HEK293 cells or (**D**), γ-irradiated (2.5 Gy) WT, *cGAS-/-* and *Sting^−^/-* BMDMos.

**Figure 5.**
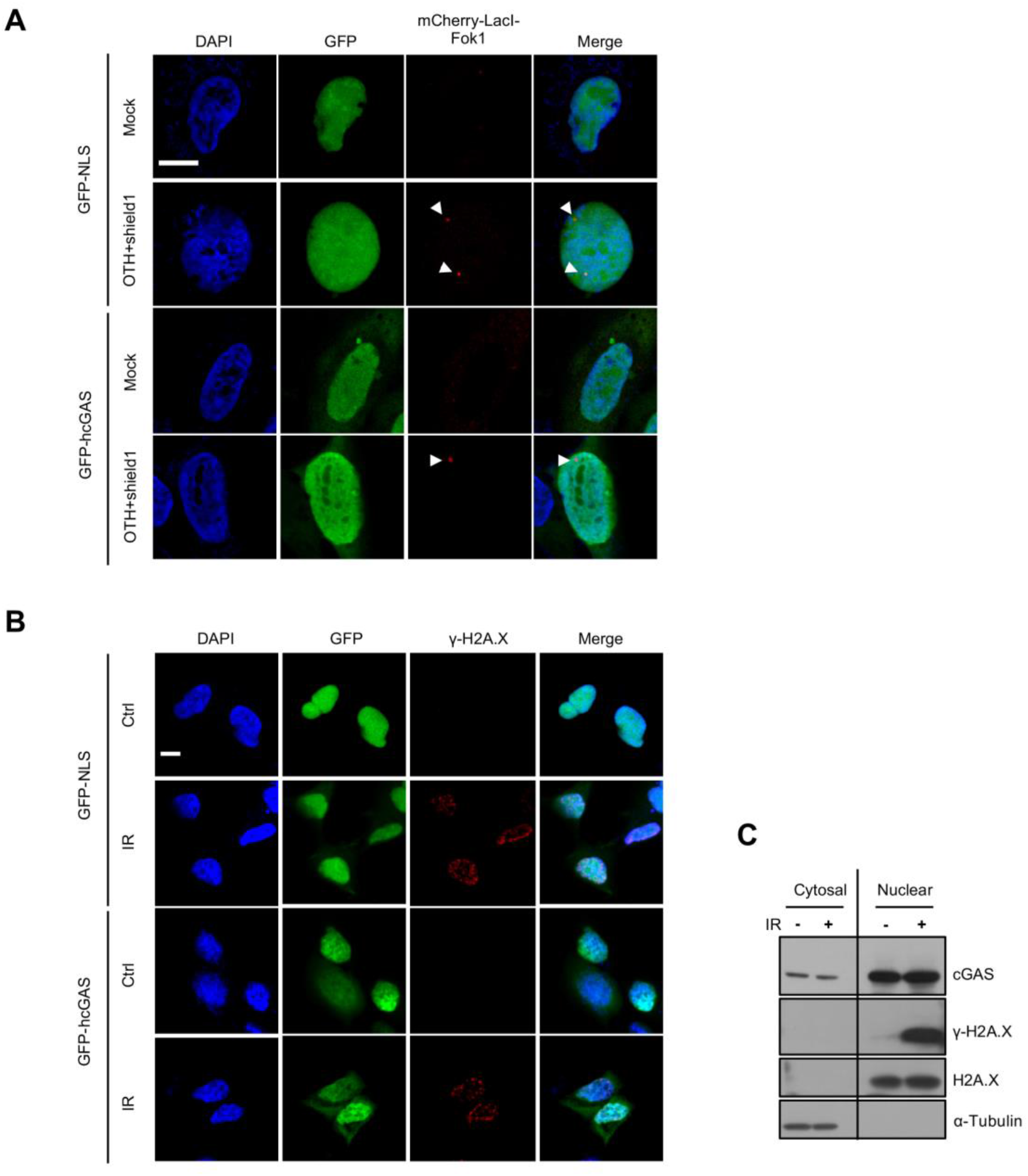
cGAS is not recruited to DSB sites. (**A**), Fluorescence images of GFP-NLS- or GFP-hcGAS-expressing U2OS-DSB reporter cells that were incubated (or not) with Shield-1 and 4-OHT to induce the expression and translocation of mCherry-LacI-FokI (red) to specific DSB sites. (**B**), Fluorescence images of GFP-NLS- or GFP-hcGAS-expressing HEK293 cells exposed (or not) to γ-irradiation (IR: 10 Gy) then stained for γ-H2A-X. (**C**), Cytosolic and nuclear pools of cGAS remain unaltered upon γ-irradiation. Immunoblots of cGAS, γ-H2A-X, H2A and Tubulin in cytosolic and nuclear fractions of γ-irradiated (10 Gy) BMDMos.

**Figure S6.**
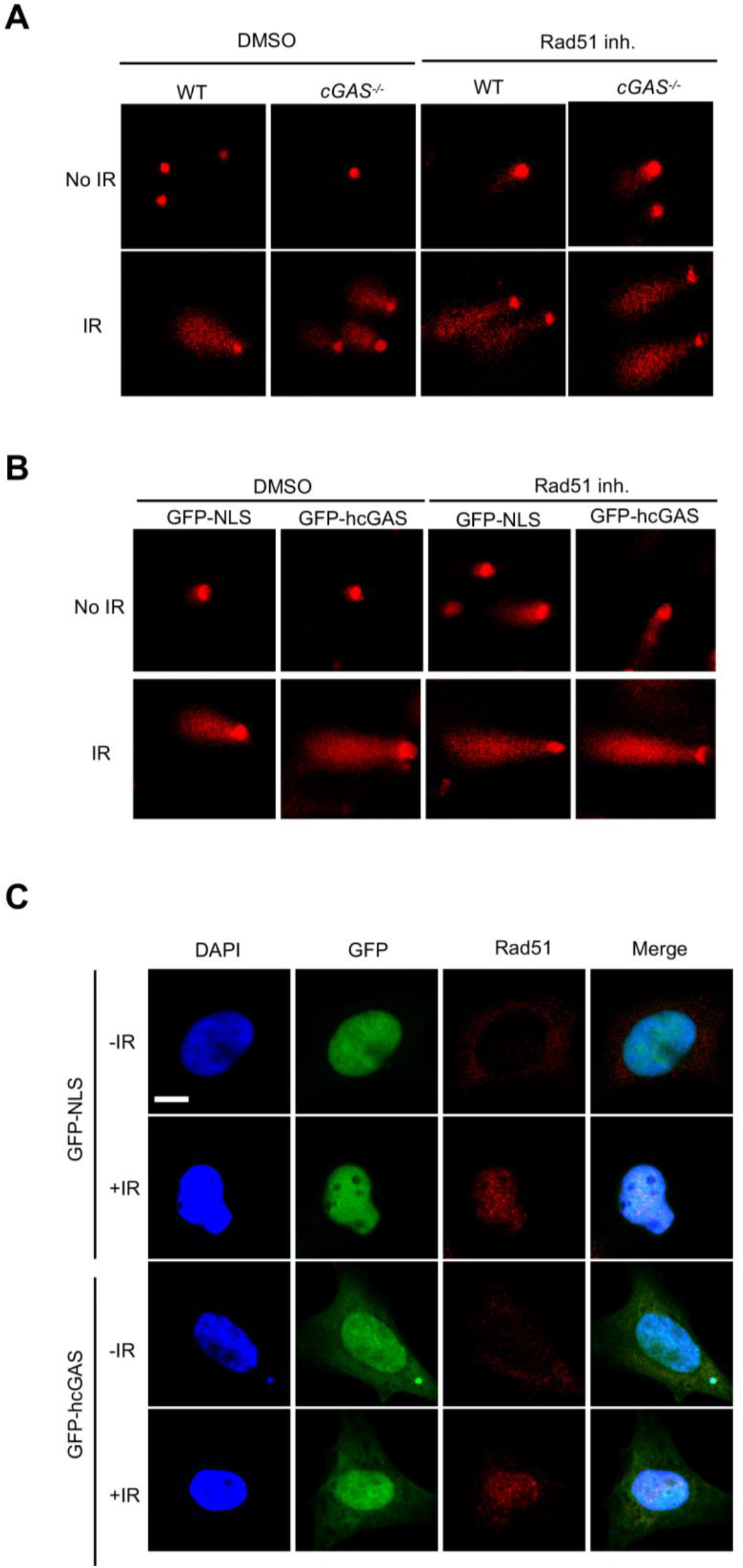
cGAS suppresses DNA repair in a RAD51-dependent manner but does not colocalize with or impede RAD51 foci formation. (**A-B**), RAD51 inhibition abrogates cGAS-mediated suppression of DNA repair. (**A**) Representative images of comet tail in (**A**) WT and *cGAS^−/−^* BMDMos or (**B**) GFP-NLS-, GFP-hcGAS-expressing HEK293 cells pretreated with the RAD51 inhibitor B02 (20 μM) 12 hour prior to γ-irradiation (10 Gy). (**C**) γ-irradiated GFP-NLS-, GFP-hcGAS-expressing HEK293 cells) stained for RAD51 (red).

**Figure S7.**
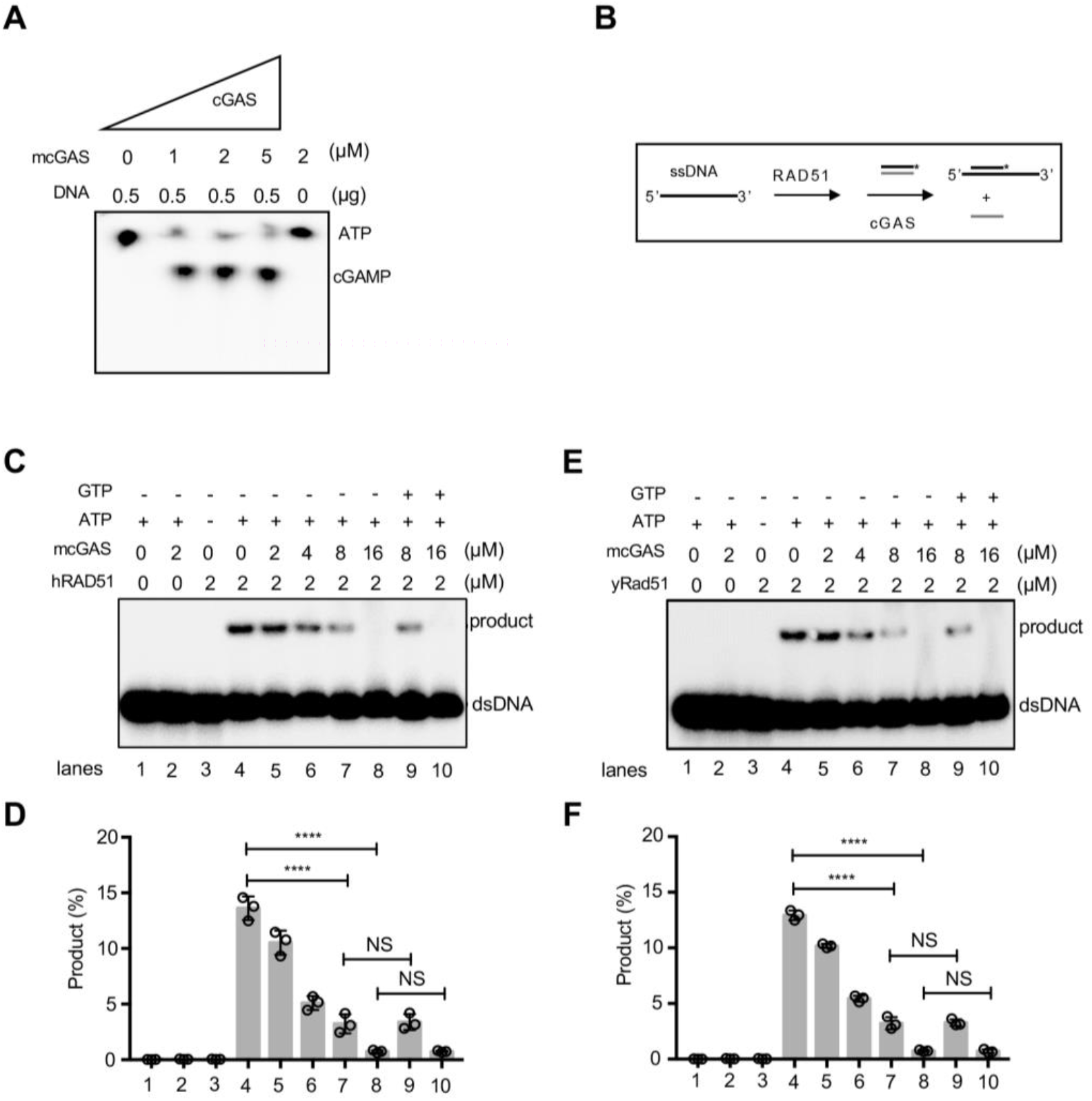
cGAS inhibits RAD51-mediated DNA strand exchange. (**A**), cGAMP synthase activity of purified mouse cGAS. (**B-F**) cGAS inhibits RAD51-mediated DNA strand exchange. Schematics of the DNA strand exchange reaction (**B**). Pre-incubation of dsDNA with cGAS protein inhibited the DNA strand exchange activity of human RAD51 (**C, D**) and yeast Rad51 (**E, F**) regardless the precursors (ATP+GTP) of cGAMP were present or not. The percentage of DNA strand exchange in each reaction was graphed as the average of triplicates ± SD., unpaired two-tailed Student’s t-test. NS: P >0.05, ****P ≤ 0.0001.

**Figure S8.**
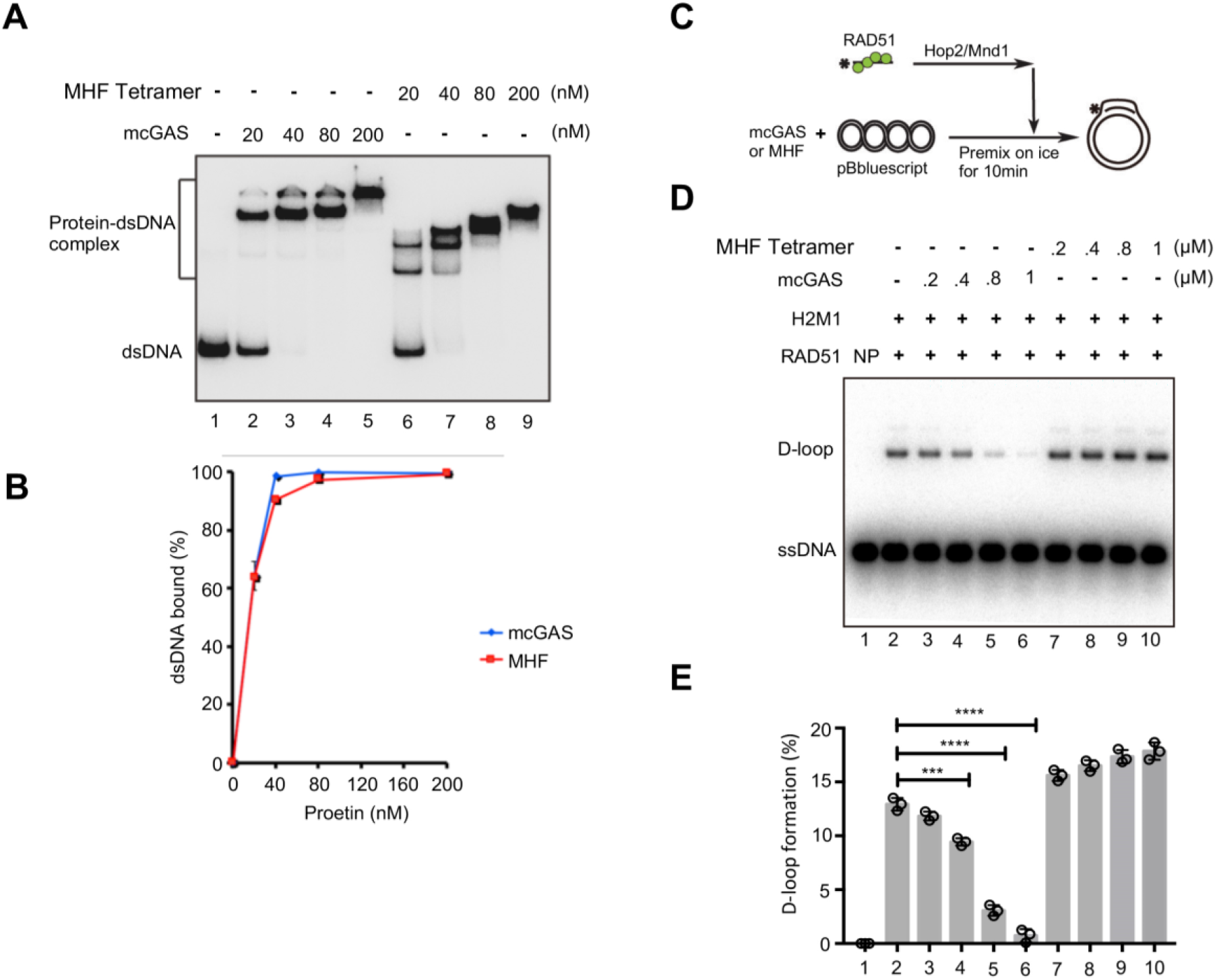
Inhibition of RAD51-mediated D-loop formation is not a ubiquitous feature of all dsDNA binding proteins. **a, b**, cGAS and MHF binds to dsDNA with similar affinity. 20-200 nM of cGAS or MHF were incubated with dsDNA at 37°C and protein-dsDNA complexes analyzed by gel separation (**a**) densitometric quantification were graphed as the average of triplicates ± SD (**b**). **c**, Schematics of the D-loop assay. **d**, Pre-incubation of template dsDNA with cGAS blocks subsequent D-loop by RAD51 while pre-incubation with the DNA binding protein MHF does not. **e**, The percentage of D-loop formed in each reaction (**d**) was graphed as the average of triplicates ± SD., unpaired two-tailed Student’s t-test. ****P ≤ 0.0001.

